# Introduction of a terminal electron sink in chloroplasts decreases leaf cell expansion associated to higher proteasome activity and lower endoreduplication

**DOI:** 10.1101/2022.07.25.501432

**Authors:** Rocío C. Arce, Martín L. Mayta, Michael Melzer, Mohammad-Reza Hajirezaei, Anabella F. Lodeyro, Néstor Carrillo

**Affiliations:** Instituto de Biología Molecular y Celular de Rosario (IBR-UNR/CONICET), Facultad de Ciencias Bioquímicas y Farmacéuticas, Universidad Nacional de Rosario (UNR), 2000 Rosario, Argentina; Leibniz Institute of Plant Genetics and Crop Plant Research, OT Gatersleben, Corrensstrasse, D-06466 Stadt Seeland, Germany

**Keywords:** electron transport, endoreduplication, flavodoxin, leaf cell expansion, leaf development, leaf size, photosynthesis, proteomics, proteasome, reactive oxygen species.

## Abstract

Foliar development involves successive phases of cell proliferation and expansion that determine the final leaf size, and is characterized by an early burst of reactive oxygen species (ROS) generated in the photosynthetic electron transport chain (PETC). Introduction of the alternative PETC acceptor flavodoxin in tobacco chloroplasts led to a reduction in leaf size associated to lower cell expansion, without affecting cell numbers per leaf. Proteomic analysis showed that components of the light-harvesting systems accumulated before electron-transport proteins, suggesting a mechanism for the early oxidative event. Flavodoxin expression did not affect biogenesis of the PETC but prevented ROS build-up through its function as electron sink. Mature leaves from flavodoxin-expressing plants were shown to contain higher levels of transcripts encoding components of the proteasome, a key negative modulator of organ size. Proteome profiling revealed that this differential accumulation initiated during expansion and led to increased proteasomal activity, whereas a proteasome inhibitor reverted the flavodoxin-dependent size phenotype. Cells expressing plastid-targeted flavodoxin displayed lower endoreduplication, also associated to decreased organ size. These results provide novel insights into the regulation of leaf growth by chloroplast-generated redox signals, and highlight the potential of alternative electron shuttles to investigate novel link(s) between photosynthesis and plant development.

**Highlight:** Modification of chloroplast redox status by expression of the cyanobacterial alternative electron sink flavodoxin decreased leaf cell expansion, which was associated with higher proteasome activity and lower endoreduplication.

## Introduction

Oxygenic photosynthesis contributes most of the global enthalpy in living systems and determines the composition of the atmosphere (Nelson, 2013). At a planetary scale, about 50% of this essential process takes place in leaves, which are thus at the heart of plant growth, seed production and ultimately agriculture. Leaf size, in particular, defines the maximal photosynthetic area and is a key factor for plant performance within a given species (Ren *et al*., 2019). After originating as lateral outgrowths of shoot apical meristems, foliar development goes through two stages: an initial phase characterized by successive cell divisions, during which specific structures such as trichomes and stomata begin to form, followed by post-mitotic cell expansion, with the transitional front extending from the leaf tip in a basipetal direction. Since dividing cells also expand, the term proliferation is preferred to describe the first phase (Traas and Monéger, 2010; Gonzalez *et al*., 2010, 2012; Andriankaja *et al*., 2012; Kalve *et al*., 2014).

Given its importance, foliar development has been the subject of intense research, revealing a complex and plastic process whose outcome in terms of size and shape depends on genetic background, leaf position and environmental conditions (Andriankaja *et al*., 2012). Both cell proliferation and expansion are controlled by the interaction of various endogenous factors such as phytohormones, reactive oxygen species (ROS), sugars and other signals, which in turn affect the expression of a multitude of genes (Traas and Monéger, 2010; Bar and Ori, 2014). Despite the wealth of information gathered, final leaf size determination remains fascinating in several ways. The size and shape are so uniform for a given species in a fixed environment that these morphological features have been widely used for taxonomic classification. However, the mutation of a single gene can dramatically change these characteristics (Barkoulas *et al*., 2008; Kalve *et al*., 2014).

Chloroplasts play a central role in plant growth, not only by providing energy and nutrients via photosynthesis but also by generating and transmitting redox-based signals (Foyer *et al*., 2017; Foyer and Hanke, 2022). For instance, transient ROS bursts accompanying both initial leaf development and transition into senescence have been traced to the photosynthetic electron transport chain (PETC), and attributed to the incomplete assembly and progressive dismantling of the chain system, respectively, which would favor adventitious energy and electron transfer to molecular oxygen. Indeed, ROS build-up shows a strong negative correlation with photosynthetic activity (Juvany et al., 2013; Muñoz and Munné-Bosch, 2018), although the specific mechanisms responsible for the oxidative bursts are not yet clear. In this sense, manipulation of photosynthetic electron transport through the incorporation of alternative electron carriers and sinks might provide a tool to understand (and handle) retrograde signals generated during photosynthesis that influence leaf development (Dietz *et al*., 2016).

Cyanobacterial flavodoxin (Fld) provides a remarkable example of this approach. Fld is a low-potential FMN-containing electron carrier, which mediates essentially the same reactions as the iron-sulfur protein ferredoxin (Fd), including electron shuttling from PSI at the PETC to a plethora of metabolic, dissipative and regulatory processes (Lodeyro *et al*., 2012; Pierella-Karlusich *et al*., 2014). In cyanobacteria and oceanic algae, Fld expression is induced by most environmental adversities to take over the activities of stress-sensitive Fd, thus allowing growth and reproduction under the adverse conditions (Pierella-Karlusich *et al*., 2014). Fld-encoding genes are absent from plant genomes (Pierella-Karlusich *et al*., 2015), but introduction of a plastid-targeted Fld into transgenic lines resulted in increased tolerance to various biotic and abiotic stresses by relieving the excess of excitation energy on the PETC and limiting ROS propagation (Tognetti *et al*., 2006, 2007; Zurbriggen *et al*., 2008, 2009; Coba de la Peña *et al*., 2010; Li *et al*., 2017; Rossi *et al*., 2017).

Noteworthy, the incorporation of Fld as an additional chloroplast redox player led to a significant transcriptional reprogramming and elicited phenotypic effects even in plants grown under normal conditions, in which Fd and Fld accumulate to similar levels (Tognetti *et al*., 2006; Ceccoli *et al*., 2012). About 1,000-1,500 leaf transcripts changed their expression patterns in response to Fld presence in plastids, as documented for both tobacco (Pierella-Karlusich *et al*., 2017) and potato (Pierella-Karlusich *et al*., 2020). Fld-expressing plants displayed higher pigment contents and photosynthetic activity per leaf cross-section (Tognetti *et al*., 2006; Ceccoli *et al*., 2012; Rossi *et al*., 2017), accompanied by an increased oxidation state of the plastoquinone pool under normal growth conditions (Gómez *et al*., 2020). They also exhibited delayed senescence and a functional stay-green phenotype, correlating with suppression of the late ROS burst (Mayta *et al*., 2018). Finally, Fld expression has been reported to decrease plant and leaf size in genetically distant species such as creeping bentgrass (Li *et al*., 2017), Arabidopsis (Su *et al*., 2018), and tomato (Mayta *et al*., 2019).

These results suggest that Fld can be used as a tool to investigate the relationship between the oxido-reductive state of the chloroplast and leaf growth, by altering the redox poise of the PETC in a tailored manner and in full illumination. Here we describe the effects of Fld presence on leaf development using a combination of morphometric, molecular and proteomic assays. We show that leaves from Fld-expressing tobacco plants grown under controlled conditions were significantly smaller than their wild-type (WT) counterparts. The reduction in size was caused by decreased cell expansion, without changes in cell proliferation or the timing of the various transitions. Proteomic analysis indicated that the biogenesis of the photosynthetic machinery was gradual, with the accumulation of light-harvesting proteins preceding that of electron transport or ATP synthesis components, which might explain the increased energy and electron transfer to oxygen and ROS build-up at this stage. While the progression of PETC biogenesis was not affected by Fld presence, the flavoprotein did suppress the early ROS burst through its function as electron sink. The effect of Fld on organ size was associated to an increase in proteasomal activity at the expansion phase, and a lower degree of endoreduplication. The possible contribution of chloroplast redox signals to the course of leaf development is discussed in the light of these observations.

## Materials and methods

### Plant material and growth conditions

Wild-type, *pfld* (for ***p***lastid-targeted ***Fld***) and *cfld* (for ***c***ytosolic ***Fld***) tobacco plants (*Nicotiana tabacum* cv. Petit Havana) were grown autotrophically in soil at 200 μmol photons m^-2^ s^-1^ with a 16-h photoperiod, 28°C/20°C, and a relative humidity of 60-80% (chamber conditions). The first true leaf emerging after the cotyledons was termed leaf 1, and leaves emerging after were numbered accordingly. The design of homozygous *pfld* and *cfld* plants has been described elsewhere (Tognetti *et al*., 2006).

### Kinematic leaf growth analysis

For leaf 1 area measurements, individual leaves (8-12 per genotype) were harvested at different days post-germination (dpg), and immediately incubated in 96% (v/v) ethanol until complete pigment removal (∼12 h). Leaf tissue was then clarified by incubation in 85% (w/v) lactic acid for 1 week. Leaves were mounted in lactic acid on glass slides, photographed at low magnification (0.75×) under a dissecting microscope (Leica MZFLIII, Wetzlar, Germany), and their areas measured with the ImageJ software (Rasband, 1997–2008, http://rsb.info.nih.gov/ij/). For leaf 10 analysis (tenth true leaf emerging after the cotyledons), 12-14 plants of each genotype were grown in 3-L pots until leaf 10 became visible in the shoot apical meristem (∼44 dpg). Leaf contour was manually drawn in a paper sheet at different dpg during development and the foliar areas were obtained from the drawn leaf contour using ImageJ. Leaf *Chl* and carotenoids contents were determined spectrophotometrically after extraction with 96% (v/v) ethanol (Lichtenthaler, 1987).

### Structural analysis of leaf tissue by light and transmission electron microscopy

To determine cell size and number, leaves 1 of at least 8 independent plants per genotype were harvested at different developmental stages and cleared as indicated above. Images of leaf cells from the adaxial palisade parenchyma layer or the adaxial epidermis were acquired through differential interference contrast microscopy (Olympus BH2, Tokyo, Japan), and their areas quantified using ImageJ. At least 100 cells were measured in 4 different regions (Fig. 1D) to calculate mean cell areas. Leaf blade area was divided by the average cell area to estimate the total number of cells per leaf. The stomatal index (SI) was calculated as: SI = [number of stomata/total number of cells (epidermal + stomata)] × 100, and averaged from 4 independent image fields of the adaxial epidermis per plant. Chloroplast coverage was defined as the ratio of total chloroplast plan area per cell plan area as determined with ImageJ in palisade parenchyma cells.

**Fig. 1.**
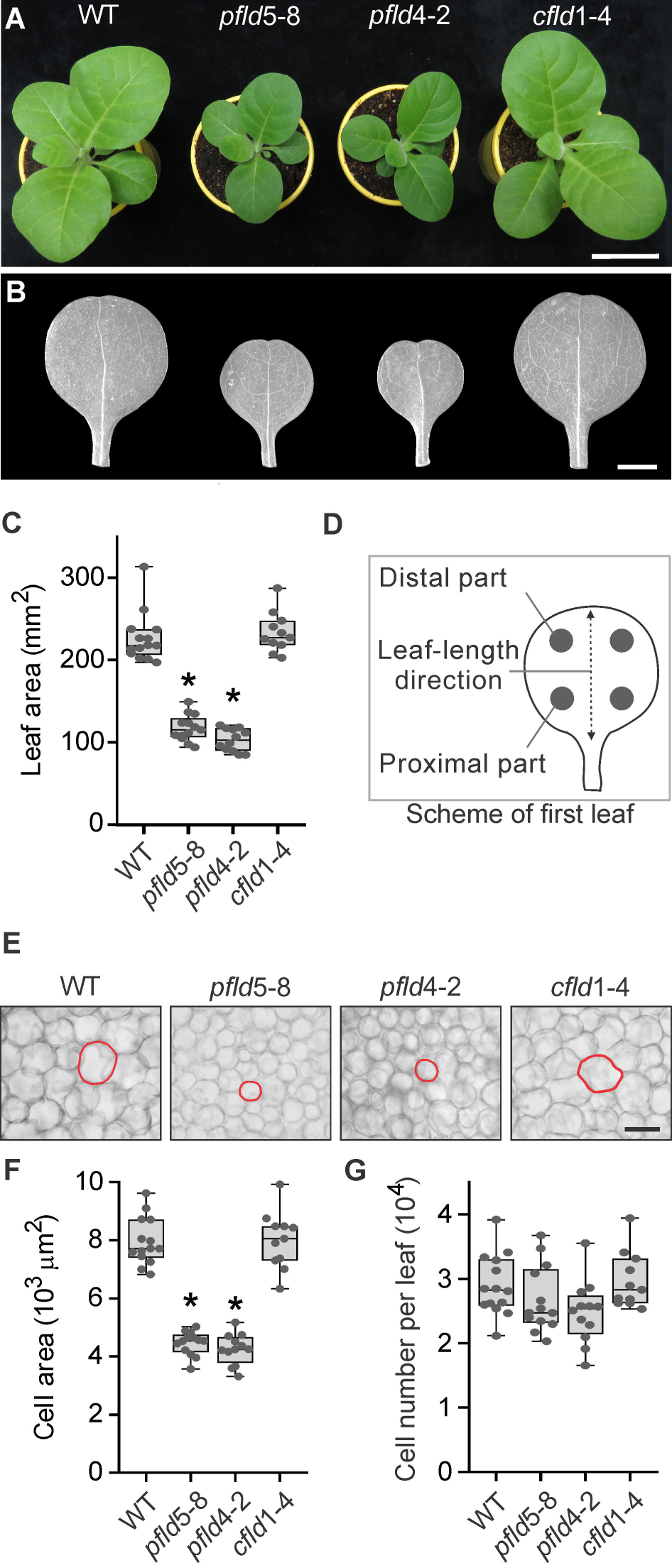
Effect of chloroplast-targeted Fld on tobacco leaf size. (A) Phenotypes of typical WT and Fld-expressing tobacco plants grown under chamber conditions at 35 dpg. Bar = 5 cm. (B) Phenotypes of leaf 1 from the same lines. Bar = 0.5 cm. (C) Areas of leaf 1 at 35 dpg determined by image analysis. (D) Leaf scheme showing where cell size was determined. Four regions of the leaf blade were used and their values averaged. (E) Representative micrographs from palisade parenchyma cells. The contour of typical cells is shown in red. Bar = 100 µm. (F) Cell areas and (G) cell numbers were calculated from clarified leaf tissue, as indicated in Materials and methods. Data are presented as box plots of 11-14 biological replicates per genotype. Central horizontal lines show the medians; box limits indicate the 25^th^ and 75^th^ percentiles; whiskers extend 1.5 times the interquartile range from the 25^th^ and 75^th^ percentiles; grey filled circles represent data points. Asterisks indicate statistically significant differences (*P <* 0.05) determined using one-way ANOVA and Tukey multiple comparison test or Kruskall–Wallis one-way ANOVA and Dunn’s multiple range test.

For histological and ultrastructural examination, leaf cuttings (2 mm^2^ in size) from the central part of leaf 1 from 4 different WT and *pfld* plants at different dpg were used for conventional and microwave-assisted fixation, substitution, resin embedding, sectioning, and microscopic analyses as described (Kraner *et al*., 2017).

### Chloroplast isolation

Intact chloroplasts were isolated from WT and *pfld*5-8 leaf 1 at 25 dpg using Percoll gradient centrifugation (Napier and Barnes, 1995). Chloroplasts were suspended in 50 mM HEPES-KOH, pH 8.0, 330 mM sorbitol, and counted after appropriate dilutions using a Neubauer chamber, an Olympus BH2 microscope, and ImageJ analysis. *Chl* contents were measured according to Napier and Barnes (1995). The average amounts of *Chl* per chloroplast were calculated by dividing the *Chl* contents per μl of chloroplast suspension by the chloroplast number per μl in the same suspension. Chloroplast numbers per cell were then estimated as the ratio between *Chl* content per cell and *Chl* content per chloroplast.

### Determination of photosynthetic parameters

While developmental changes were followed in leaf 1 throughout the manuscript, small size precluded its use for photosynthetic assays, which require a minimal measuring leaf area. Leaf 10 was used instead, after confirming that its developmental pathway was similar to that of leaf 1 (see Results), as its size in the proliferative phase was large enough for photosynthetic determinations, as opposed to other leaves of the same developmental stage emerging before. *Chl a* fluorescence and P700 absorbance were measured simultaneously at 25°C using a Dual-PAM-100 system (Heinz Walz, Effeltrich, Germany). In all cases, plants were dark-adapted for 3 h to inactivate the Calvin-Benson cycle, and measurements were carried out at 200 μmol photons m^-2^ s^-1^. Light curves were determined from 35 to 2000 μmol photons m^-2^s^-1^. Parameters related to PSI and PSII photochemical responses were directly obtained from the Dual-PAM-100 software (Baker, 2008). Light-dependent CO_2_ uptake rates were measured on a LI-6400 portable photosynthesis system (LI-COR, Lincoln, NE). The CO_2_ concentration of the air entering the leaf chamber and the temperature were adjusted to 400 ppm and 25°C, respectively. Photosynthetic photon flux densities ranging from 0 to 1500 μmol photons m^-2^s^-1^ were supplied by a controlled halogen light source.

### ROS determination

ROS levels were measured in leaf 10 to facilitate direct comparison with photosynthetic activities. A modified version of the FOX II assay (DeLong *et al*., 2002) was used to quantify peroxides in cleared leaf extracts throughout leaf development. Two-hundred mg of leaf tissue were ground in liquid nitrogen and extracted with 300 µl of an 80:20 ethanol/water mixture, containing 0.01% (w/v) butylated hydroxytoluene. After centrifugation at 3,000 *g* for 10 min, 30 µl of the supernatant were mixed with 30 µl of 10 mM Tris–phenyl phosphine (a peroxide reducing agent), or the same volume of methanol. The mixtures were incubated for 30 min at 25°C, 100 µl of FOX reagent (100 µM xylenol orange, 4 mM butylated hydroxytoluene, 250 µM ferrous ammonium sulphate, 25 mM H_2_SO_4_ in 90% methanol) were then added to each sample, and the absorbance at 560 nm was recorded 10 min after reagent addition. The absorbance differences between equivalent samples with or without Tris–phenyl phosphine indicate the levels of peroxides, which were calculated using an H_2_O_2_ standard curve.

### RNA isolation and quantitative real-time PCR analysis

Total RNA was isolated from 100 mg of tobacco leaf tissue using the TriPure reagent (Roche Sigma-Aldrich, St. Louis, MO), according to the manufacturer’s instructions, and reverse-transcribed with the MMLV enzyme (Invitrogen, Carlsbad, CA) and oligo(dT)_12-18_. Real-time PCR reactions were carried out in a Mastercycler^®^ ep*realplex*^4^ thermocycler (Eppendorf, Hamburg, Germany) employing Platinum Taq DNA polymerase (Invitrogen), and SYBR Green I (Roche Sigma-Aldrich) to monitor the synthesis of double-stranded DNA. Relative transcript levels were determined for each sample and normalized against the levels of tobacco elongation factor 1α (EF1α) cDNA (Schmidt and Delaney, 2010). Primers (Invitrogen) were designed using the “Primer3Plus” software with an annealing temperature of 55°C (Table S1).

### Protein extraction for proteomic analysis

Four biological replicates of leaf 1 were collected at 6, 16 and 25 dpg and ground individually in liquid nitrogen. The resulting powder was suspended in extraction buffer containing 50 mM Tris-Cl pH 8, 0.5% (w/v) SDS, 1 mM phenyl methyl sulfonyl fluoride, 10% (v/v) glycerol, 1 mM DTT, 5 mM MgCl_2_ and 1 mM EDTA, in a 2:1 buffer:tissue ratio, and then vortexed thoroughly. After centrifugation at 7,500 *g* and 4°C for 10 min, the supernatant was separated and the protein concentration was determined according to Bradford (1976).An adequate volume of loading buffer (50 mM Tris-Cl pH 6.8, 2% SDS, 4% glycerol, 0.11% (w/v) bromophenol blue, 0.1 M DTT) was added and the samples were kept at −80°C until use. Protein extracts were subjected to SDS-PAGE on 12% (w/v) polyacrylamide gels and digested with trypsin according to Link and Labaer (2009).

### LC-MS analysis

Peptide separations were carried out on a nanoHPLC Ultimate3000 (Thermo Scientific, Waltham, MA) using an EASY-Spray ES903 nano column (50 cm × 50 μm ID, PepMap RSLC C18). Samples (3.5 µl) were injected into the column equilibrated with 4% solvent B (0.1% formic acid in acetonitrile) in solvent A (0.1% formic acid in water). The flow rate of the mobile phase was 400 nl min^-1^ using different fractions of solvent B in solvent A. The gradient profile was set as follows: 4-30% solvent B for 114 min, 30-80% solvent B for 14 min and 80% solvent B for 2 min. MS analysis was performed using a Q-Exactive HF mass spectrometer (Thermo Scientific). For ionization, 1.9 kV of liquid junction voltage and 250°C of capillary temperature were used. The full scan method employed a mass selection of an m/z 375–2,000, an Orbitrap resolution of 120,000 (at m/z 200), a target automatic gain control value of 1e6, and maximum injection times of 100 ms. After the survey scan, the 7 most intense precursor ions were selected for MS/MS fragmentation.

Fragmentation was performed with a normalized collision energy of 27 eV and MS/MS scans were acquired with a dynamic first mass. The automatic gain control target was 5e5, resolution of 30,000 (at m/z 200), intensity threshold of 1.5e5, isolation window of 1.4 m/z units and maximum injection times of 55 ms. Charge state screening was enabled to reject unassigned, singly charged, and equal or more than 6-protonated ions. A dynamic exclusion time of 15 s was used to discriminate against previously selected ions.

### MS data analysis

MS data were analyzed with MaxQuant (V: 2.1.0.00) using standardized workflows. Mass spectra *.raw files were searched against a database from *N. tabacum* (UniProt ID UP000084051). Precursor and fragment mass tolerance were set to 10 ppm and 0.02 Da, respectively, allowing 2 missed cleavages, cysteine carbamidomethylation as a fixed modification, methionine oxidation and N-terminal acetylation as variable modifications. Further downstream data processing and statistical tests were carried out with the Perseus software (V: 1.6.15.0; Tyanova *et al*., 2016). Pairwise comparisons were performed with a permutation-based false discovery rate threshold of 0.05. The differentially expressed proteins (DEPs) with fold change (FC) > 1 at *P* < 0.05 were considered to be proteins with significantly up-regulated expression, while DEPs with FC < -1 and *P* < 0.05 were determined to be proteins with significant down-regulation. DEPs were annotated according to the Kyoto Encyclopedia of Genes and Genomes (KEGG) identifiers (https://www.kegg.jp/), and the KEGG pathway graphs were generated in R language (http://www.r-project.org/) using the *Pathview* package (Luo and Brouwer, 2013). To identify relative abundance patterns in proteins related to the proteasome system, a hierarchical clustering of normalization data was performed using the expression function within Heatmapper (Babicki *et al*., 2016), with the “Average Linkage” clustering method and the “Euclidean” distance measurement method.

### Proteasome activity and inhibition assays

Proteasome activity was measured by spectrofluorometry using the 7-amino-4-methylcoumarin (AMC)-labeled fluorogenic substrate succinate-LLVY-AMC (Roche Sigma-Aldrich) in cleared extracts of leaf 1 at different dpg (Üstün and Börnke, 2017). Briefly, 100 mg of leaf tissue were frozen in liquid nitrogen and ground in 200 µl of extraction buffer (50 mM HEPES-KOH pH 7.2, 2 mM ATP, 2 mM DTT, 250 mM sucrose). After centrifugation, the protein concentration of the supernatant, as determined by the Bradford (1976) procedure, was adjusted to 1 µg µl^−1^ with extraction buffer. Fifty µl of the protein solution were mixed with 220 µl of proteolysis buffer (100 mM HEPES-KOH pH 7.8, 5 mM MgCl_2_, 10 mM KCl, 2 mM ATP). The reaction was started after 5 min at 30°C by the addition of 0.2 mM Suc-LLVY-AMC. Released AMC was measured every 2 min using a Synergy HTX spectrofluorometer (Biotek, Winooski, VT) with excitation at 360 nm and emission at 460 nm.

For proteasome inhibition assays, seedlings were grown on 0.8% (w/v) agar containing half-strength Murashige and Skoog (1962) medium (0.5 × MS-agar) until 8 dpg. Subsequently, the aerial parts were dipped for 30 s in 1 ml of 0.01% (v/v) Tween-20 aqueous solution containing 50 µM MG132 (Roche Sigma-Aldrich) in DMSO, or the equivalent volume of DMSO (control), and the plants were cultured in 0.5 × MS-agar for 7 more days. Shoots were photographed, excised and their fresh weight determined.

### Estimation of DNA endoreduplication

Nuclei extraction and population measurements were carried out as described by Fina *et al*. (2017), with minor modifications. Briefly, ten leaves 1 from individual plants of each genotype at 25 dpg, with their central vein removed, were cut in ∼1-mm slices with a razor blade in a petri dish containing 1 ml of a pre-chilled (4°C) medium made up of 20 mM MOPS pH 7.0, 30 mM sodium citrate, 45 mM MgCl_2_, and 1% (v/v) Triton X-100. After gently pipetting the solutions to obtain homogenates, they were filtered through a 100-μm mesh, supplemented with 80 μl of 0.2 mg ml^-1^ propidium iodide, 50 mg ml^-1^ RNase, incubated for 30 min, and finally read through a Cell Sorter BD FACSAria II flow cytometer (BD Biosciences, San Jose, CA). The endoreduplication index (EI) was calculated from the percentages of each ploidy class with the formula: EI = [(0 × %2C) +(1 × %4C) +(2 × %8C) + (3 × %16C) +(4 × %32C)]/100. Experiments were carried out in triplicate, and at least 5,000 nuclei were analysed each time.

### Statistical analyses

Data were analysed using one-way ANOVA and Tukey or Duncan multiple range tests. When the normality and/or equal variance assumptions were not met, Kruskall–Wallis one-way ANOVA between lines, and Dunn’s multiple range tests were used. Significant differences refer to statistical significance at *P* < 0.05.

## Results

### Fld expression in chloroplasts of tobacco plants decreases the rate of leaf cell expansion

WT and Fld-expressing plants were grown under chamber conditions (Materials and methods) to isolate the effects of the flavoprotein on leaf development from those related to stress protection that might also affect organ size. Independent tobacco lines *pfld*5-8 and *pfld*4-2 contain similar levels of Fld in chloroplasts and display an equivalent degree of stress tolerance (Tognetti *et al*., 2006). Typical phenotypes of WT and *pfld* plants at 35 dpg showed that Fld expression in plastids led to a significant decrease in leaf size in the absence of environmental pressures (Fig. 1A-C). Size reduction was observed from 12 dpg on, and affected all leaves (Supplementary Fig. S1A, B). A homozygous line expressing high levels of Fld in the cytosol (*cfld*1-4; Tognetti *et al*., 2006) exhibited a WT phenotype (Fig. 1A-C; Supplementary Fig. S1A, B), indicating that chloroplast location was mandatory for the Fld effect. These results thus concur with the size reduction reported for creeping bentgrass, Arabidopsis and tomato plants expressing a plastid-located Fld (Li *et al*., 2017; Su *et al*., 2018; Mayta *et al*., 2019).

As a first step to characterize the mechanism underlying this size reduction, different regions of leaf 1 were sampled (Fig. 1D) to estimate cell size by microscopic analysis. At 35 dpg, the average area of mesophyll cells belonging to the palisade parenchyma was decreased by ∼50% in mature *pfld* leaves compared to WT and *cfld* siblings (Fig. 1E, F), correlating with the reduction in leaf area (Fig. 1C). The total numbers of cells per leaf, instead, were not significantly modified (Fig. 1G). Epidermal pavement cells exhibited a similar pattern (Supplementary Fig. S2A-C), whereas SI values were not affected by the presence of the flavoprotein (Supplementary Fig. S2D). When the same type of experiment was carried out at 6 dpg, no significant differences in cell size were observed between lines (Supplementary Fig. S3). The results suggest that Fld-dependent changes in chloroplast redox status did not affect the early stages of leaf development associated to cell proliferation, but had a detrimental impact on cellular expansion.

To determine if the smaller cells of *pfld* leaves were the result of lower rates of cell expansion and/or shortening of the expansion phase, a kinematic analysis of leaf and cell development was carried out in WT and *pfld* plants. As shown in Fig. 2A, average areas of leaf 1 were similar for all lines up to 9 dpg, but *pfld* leaves grew significantly slower than WT counterparts after that time. Growth ceased at about 20 dpg in all lines (Fig. 2A), indicating that the onset of leaf maturity was not affected by the presence of the additional redox carrier. Leaves emerging later than leaf 1 displayed similar behavior, as illustrated for leaf 10 in Supplementary Fig. S4.

**Fig. 2.**
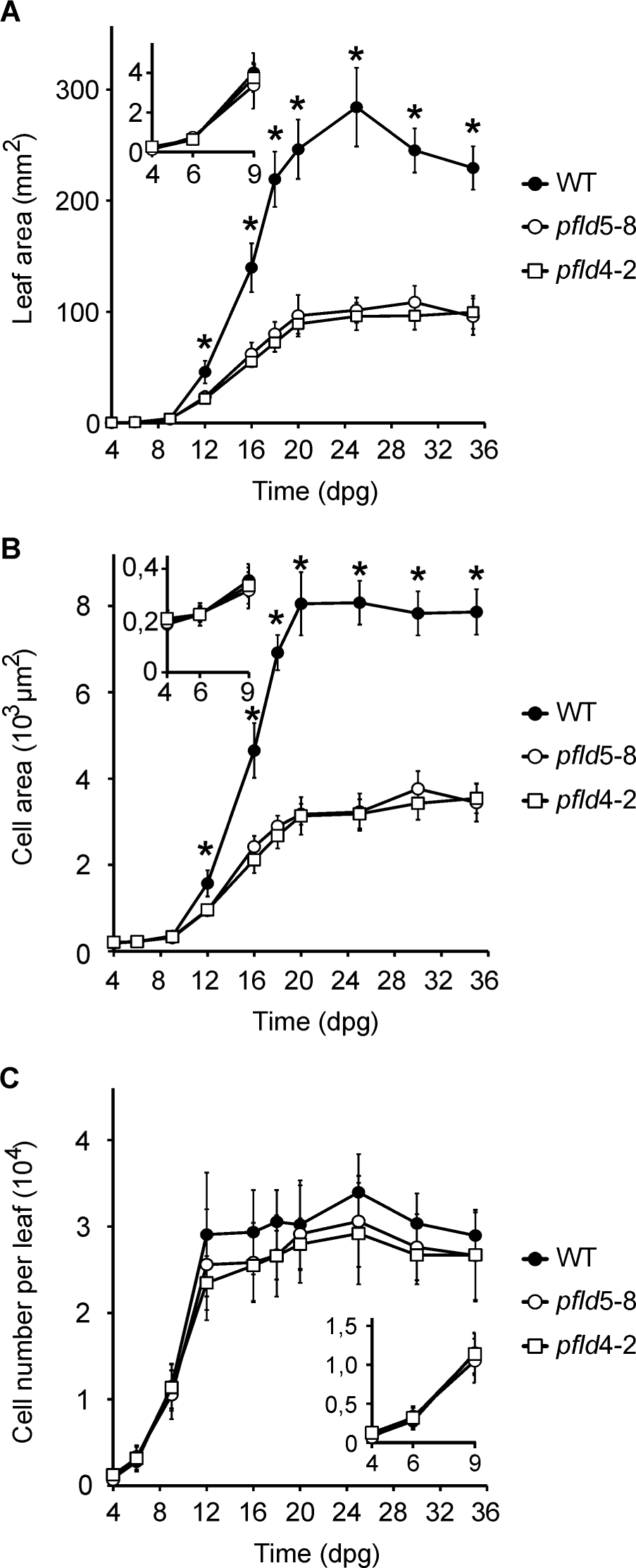
Kinematic analysis of leaf growth in wild-type and *pfld* plants. The first leaves of WT and *pfld* plants were collected at the indicated dpg for determination of leaf and cell area. (A) Changes in leaf area during development. Data are means ± SD of at least 8 independent plants per line. (B) Average areas of adaxial sub-epidermal palisade parenchyma cells during leaf development. Cell areas were determined in 4 regions as indicated in Fig. 1D. Data are represented as means ± SD (*n* = 100 cells from at least 8 leaves). (C) Changes in the number of cells per leaf during development. Cell numbers were estimated in individual leaves as described in Materials and methods. Data are means ± SD from at least 8 leaves. Insets show a portion of the data that was magnified to show the absence of differences in the early stages of leaf development. Asterisks indicate statistically significant differences between WT and both *pfld* lines (*P <* 0.05) determined using one-way ANOVA and Tukey multiple comparison test, or Kruskall–Wallis one-way ANOVA and Dunn’s multiple range test.

Cellular analysis over time showed a good correlation with these phenotypic features. Cell numbers increased at similar rates in WT and *pfld* leaves, reaching their maxima at ∼12 dpg irrespective of the genotype, whereas post-mitotic cell expansion continued up to ∼20 dpg in all lines, with *pfld* cells expanding at lower rates compared to WT siblings (Figure 2B, C). The number of cells per unit area declined steadily throughout organ development, as it grows in size, but *pfld* leaves began to pack a higher number of smaller cells per cross-section upon entering the expansion phase, stabilizing at a ∼2-fold higher cell density than WT siblings at the end of this stage (Supplementary Fig. S5). The results thus show that Fld activity specifically affected expansion rates with no appreciable effects on cell proliferation or the timing of exit from proliferation and expansion (Fig. 2B, C).

### Fld presence affects chloroplast numbers and chlorophyll contents

Previous observations have shown that *pfld* plants contained more *Chl* and carotenoids per leaf area than WT siblings (Tognetti *et al*., 2006; Rossi *et al*., 2017; Gómez *et al*., 2020). Analysis of pigment contents along the various stages of leaf development confirmed those observations (Supplementary Fig. S6). Differences between lines were modest at the proliferative stage, but the levels of *Chl a* and carotenoids increased in *pfld* leaves from the beginning of the expansion phase, well above those of their WT counterparts (Supplementary Fig. S6A, D). *Chl b* contents only began to diverge between lines at the onset of leaf maturity, when pigment levels were already declining (Supplementary Fig. S6B).

Then, Fld activity affected both cell density and pigment contents per area (Supplementary Fig. S5, S6). *Chl* and carotenoids accumulation per cell kept the pace of cell division during the entire proliferative stage, resulting in similar pigment levels per cell for all genotypes (Supplementary Fig. S7). During expansion and early leaf maturity, the increase in cell packing surpassed pigment build-up, and *pfld* leaves actually contained less *Chl* and carotenoids per cell than WT siblings (Supplementary Fig. S7). The two Fld effects cancelled each other at 25-30 dpg, and mature leaves from all lines exhibited similar pigment contents per individual cell (Supplementary Fig. S7).

Individual chloroplasts isolated from mature leaves of *pfld* plants contained 30-40% more *Chl* than WT counterparts (Supplementary Fig. S8A). However, cells from mature *pfld* leaves had less chloroplasts (Supplementary Fig. S8B), likely reflecting spatial constraints on plastid proliferation in these smaller cells. The positive correlation between plastid proliferation and cell size has been reported previously (Jarvis and López-Juez, 2013; Kawade *et al*., 2013).

Image analysis of chloroplasts from palisade parenchyma cells visualized by electron microscopy did not show consistent differences between lines in plastid ultrastructure, average area or grana stacking, as illustrated for WT and *pfld*5-8 plastids in Supplementary Fig. S9A-C. Despite their lower number and similar size, chloroplast coverage per cell was higher in *pfld* leaves (Supplementary Fig. S8C), as the reduction in cell size offset the decline in plastid numbers. Collectively, the results indicate that the increased *Chl* levels per area in mature *pfld* leaves (Supplementary Fig. S6; Tognetti *et al*., 2006; Ceccoli *et al*., 2012; Gómez *et al*., 2020) results from the combination of two factors: a higher density of smaller cells per unit of leaf cross-section (Supplementary Fig. S5), and increased *Chl* contents per chloroplast at leaf maturity (Supplementary Fig. S8A).

### Fld expression does not affect biogenesis of the photosynthetic electron transport chain but suppress ROS build-up in developing tobacco leaves

Biogenesis of the PETC is reported to proceed stepwise during leaf development (Andriankaja *et al*., 2012; Juvany *et al*., 2013; Huang *et al*., 2018). To further characterize this process, we performed a label-free quantitative proteomic analysis of WT and *pfld* leaves. According to the growth profile of leaf 1, we chose three representative developmental stages: proliferation at 6 dpg, expansion at 16 dpg, and maturity (fully expanded leaves) at 25 dpg.

A total of 8,911 individual proteins could be identified and assigned from the collected samples, comprising all genotypes and developmental phases. Comparison between stages revealed an age-dependent decline in the number of proteins detected: 7,928 at 6 dpg, 5,215 at 16 dpg, and 4,772 at 25 dpg. From this richness of data, we focused on the biogenesis of the PETC by following accumulation of chain components along leaf growth. DEPs associated with this functional category were annotated according to the KEGG pathways and plotted using Pathview (Luo and Brouwer, 2013). Results obtained with WT plants are shown in Fig. 3, and those corresponding to *pfld* lines in Supplementary Fig. S10 and S11. In this representation, a continuous decline along time indicates that accumulation of the corresponding protein already peaked at the first data sampling (6 dpg in our case).

**Fig. 3.**
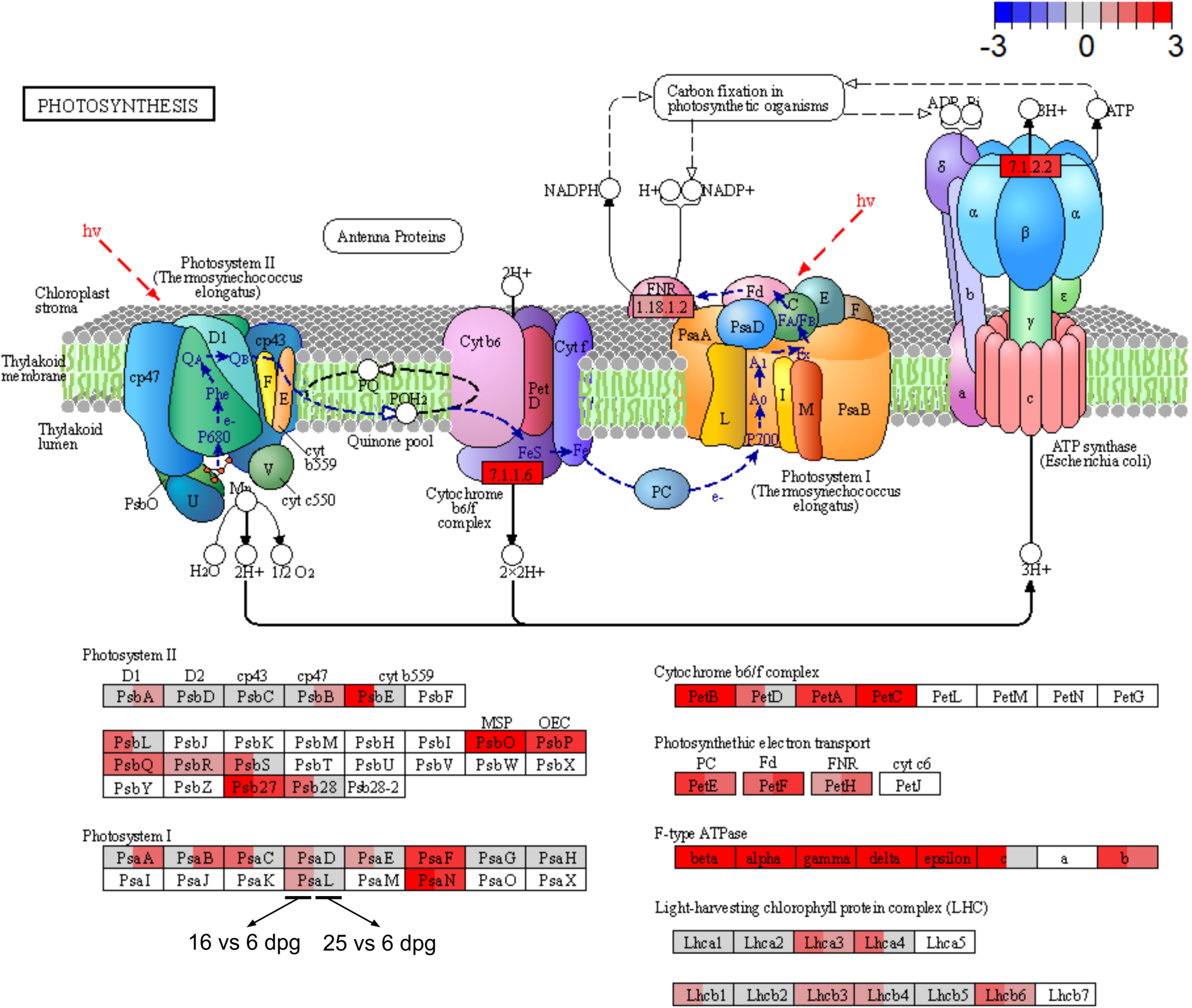
Expression of photosynthesis-related proteins through leaf development in WT tobacco leaves. KEGG pathway changes were analyzed using Pathview. The color scale indicates the extent of up-(red) or down-regulation (blue). The upper part of the figure shows a scheme of the PETC with its multiple components (each displaying the acronym of its name), whose regulation is detailed in the lower part. Rectangles in the upper panel indicate the activities of selected complexes and enzymes as defined by the IUBMB: EC 7.1.1.6, plastoquinone-plastocyanin reductase; EC 1.18.1.2, ferredoxin-NADP^+^ reductase; EC 7.1.1.2, ATP synthase. Rectangles in the lower panel show the modulation of individual subunits. Each rectangle (upper and lower panels) was divided into 2 sections, which represent proteins modulated at 16 dpg vs 6 dpg (left), and 25 dpg vs 6 dpg (right). Empty boxes correspond to components that were not detected in the proteomic analysis. Activities depicted in the upper panel are suggested to be induced at the two developmental stages according to Pathview, as indicated by the red filling. Details on the experimental design and data analysis are given in Materials and methods.

Fig. 3 shows that many components of the PETC increased their levels during growth of WT leaves. Noteworthy, build-up of most light-harvesting proteins peaked at the expansion phase (16 dpg), whereas components of PSII, the cytochrome *b*_6_*f* complex, electron shuttles, Fd-NADP(H) reductase and the proton ATP synthase increased at later stages, and only attained their maximal accumulation at leaf maturity (Fig. 3). Photosystem I exhibited an intermediate pattern, with peaks of PSI subunit accumulation distributed between 16 dpg and 25 dpg (Fig. 3). Proteomic analysis also revealed that enzymes involved in the synthesis of tetrapyrroles, the biochemical precursors of chlorophylls, exhibited their highest abundance at 6 dpg, when cells were still proliferating, and declined at later stages (Supplementary Fig. S12). Transcriptional and proteomic profiling of Arabidopsis developing leaves yielded essentially the same results (Andriankaja *et al*., 2012; Omidbakhshfard *et al*., 2021). These observations suggest that during biogenesis of the PETC, light harvesting precedes photochemistry, which might explain the early ROS burst of developing leaves (Muñoz and Munné-Bosch, 2018).

To determine how the stepwise assembly of the PETC affected photosynthetic activities, we measured rates of electron transport during leaf development. Small leaf size, especially at the early stages, precluded the use of leaf 1 for photosynthetic determinations, so leaf 10 was used instead to estimate the quantum yields of PSI and PSII (φ_PSI_ and φ_PSII_, respectively), which provide an estimation of the electron flow through these photosystems. Measurements carried out in WT plants at 44 dpg, when cells of leaf 10 are for the most part at the proliferative stage and no difference in leaf size is observed between lines (Supplementary Fig. S4), yielded significantly lower φ_PSI_ and φ_PSII_ values than those of expanding leaves (Fig. 4A, B), in agreement with the proteomics results. Quantum yields increased steadily with leaf age for both photosystems, reaching a maximum at the end of the expansion phase and then declining to initial values (Fig. 4A, B). ROS levels were determined in cleared extracts of the same WT leaves. They showed a negative correlation with electron transport rates (Fig. 4C). The results suggest that during the early stages of WT leaf growth, part of the electron flow estimated by φ_PSI_ and φ_PSII_ was used to reduce oxygen to ROS.

**Fig. 4.**
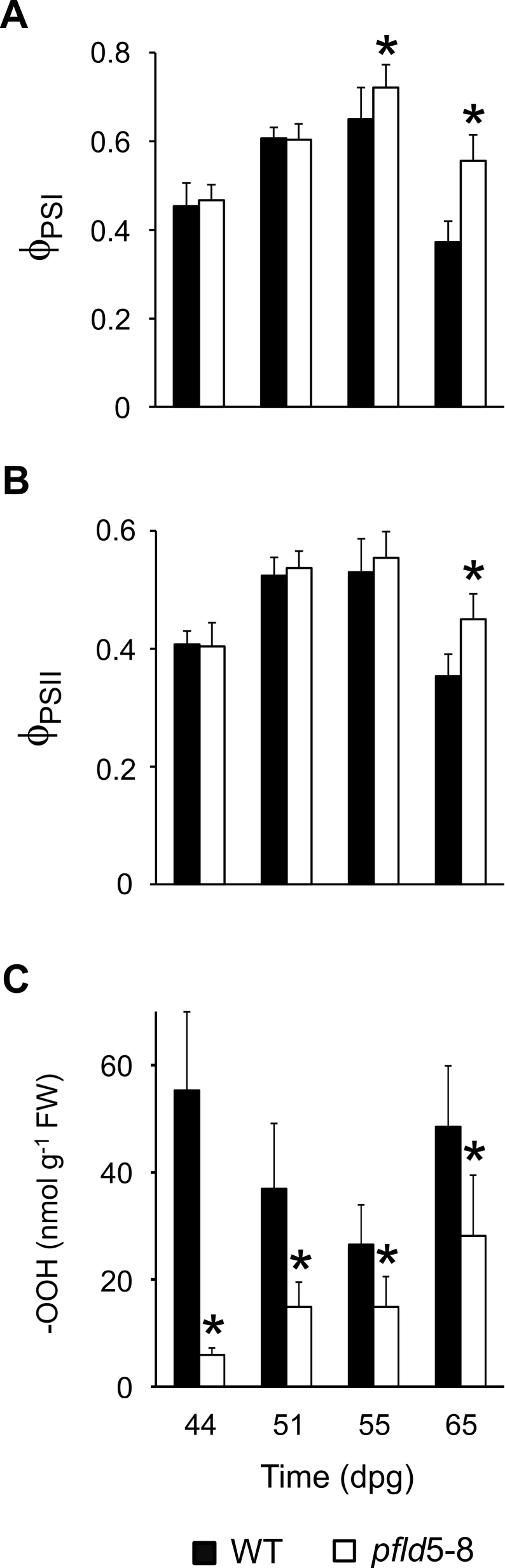
Effect of plastid Fld expression on electron transport rates and ROS production during leaf development. Determination of the quantum yields of (A) PSI and (B) PSII (__PSI_ and __PSII_, respectively) was carried out in leaf 10 (n = 5-7) at the indicated dpg. (C) Contents of total peroxides (–OOH) were measured in cleared extracts of leaf 10 (n = 3). Data reported are means ± SD, and asterisks indicate statistically significant differences (P < 0.05) determined using one-way ANOVA and Duncan’s multiple comparison test.

Chloroplast Fld did not affect the accumulation of PETC components during leaf development, nor did it improve photosynthetic activity during proliferation and early expansion of *pfld*5-8 leaves (Supplementary Fig. S10, S11; Fig. 4A, B). The decline of electron transport rates in older leaves was instead partially prevented by the plastid-targeted flavoprotein (Fig. 4A, B), in line with previous observations (Mayta *et al*., 2018). Analysis of light-response curves indicated that the differential effect of Fld on electron transport (Supplementary Fig. S13A, B) was accompanied by significant increases of CO_2_ assimilation rates in *pfld* plants (Supplementary Fig. S13C), and only became evident during late expansion for all measured parameters. It is worth noting, in this context, that no differences in electron transport or CO_2_ fixation rates were detected at the growth light intensity of 200 μmol photons m^-2^ s^-1^ (Supplementary Fig. S13). Also, and despite the lack of Fld influence on the photosynthetic activity of immature leaves, the flavoprotein did decrease ROS propagation at all stages of leaf development including the early oxidative burst (Fig. 4C), presumably through its role as electron sink.

### Fld effects on leaf size correlated with increased activity of the proteasome

Among the various factors influencing organ development, proteasome activity has been negatively correlated with leaf size (Kurepa *et al*., 2009; Sonoda *et al*., 2009), and previous observations indicated that Fld presence in chloroplasts led to generalized induction of proteasome-encoding transcripts in mature leaves of tobacco and potato plants compared to WT siblings (Pierella-Karlusich *et al*., 2017, 2020). The time dependence of this Fld-dependent up-regulation was investigated for several components of both the 20S proteasomal core and the 19S regulatory complex by quantitative real-time PCR (Supplementary Fig. S14). The results show that by early cell expansion, the levels of all transcripts assayed were already significantly higher in *pfld*5-8 leaves compared to WT siblings (Supplementary Fig. S14).

Enhanced build-up of proteasome-encoding transcripts in *pfld* leaves translated into protein levels, as revealed by proteomic analysis (Fig. 5A). Interestingly, the contribution of proteasomal proteins to the total leaf proteome of WT plants was maximal at 6 dpg, and decreased in successive stages (Supplementary Fig. S15). A similar decline in proteasome-related transcripts and proteins along leaf development was reported for Arabidopsis (Andriankaja et al., 2012; Omidbakhshfard *et al*., 2021). Fld-driven differential accumulation of proteasomal components was already evident at 16 dpg (Fig. 5A), correlating with the lower rates of cell expansion displayed by these leaves. It was particularly prominent for components of the 20S core particle, whereas build-up of subunits belonging to the regulatory complex was lower in average and protracted (Fig. 5B). Indeed, analysis of protein abundance along leaf development indicates that Fld presence preferentially prevented the age-dependent decline of proteins belonging to the proteasome core, compared to components of the regulatory particle (Supplementary Fig. S15).

**Fig. 5.**
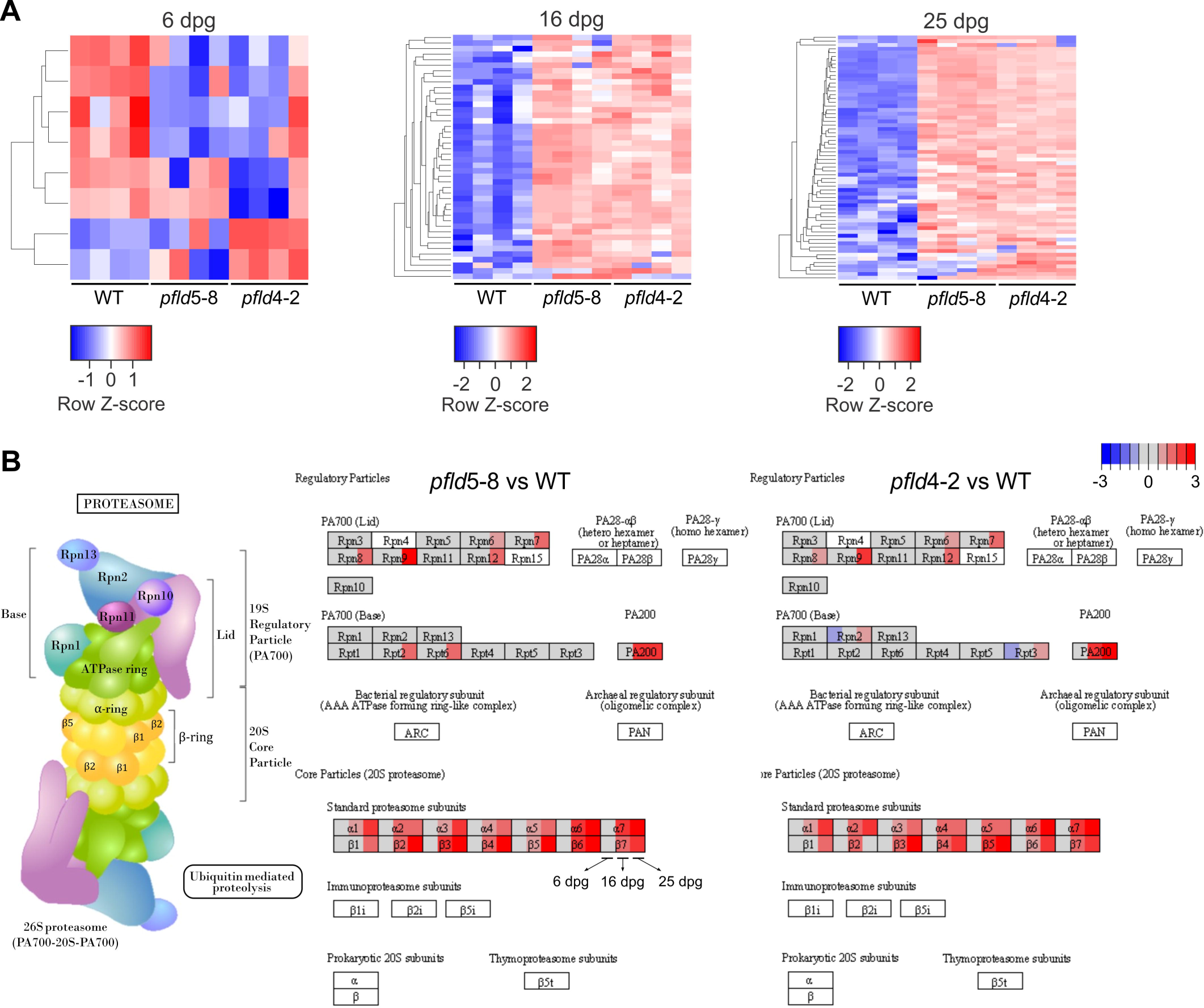
Expression of proteasome-related proteins during leaf development in WT and Fld-expressing plants. (A) Hierarchical clustering of abundance patterns for proteins related to the proteasome in leaf 1 of WT, *pfld*5-8 and *pfld*4-2 plants at 6, 16 and 25 dpg. (B) Regulation of components of the proteasome complex through development. In the visualization, each rectangle was divided into 3 subsections corresponding to 6, 16 and 25 dpg. The color scale indicates the extent of up-(red) or down-regulation (blue). Empty boxes indicate components that were not detected in the proteomic analysis. Further experimental details are given in Materials and methods.

In line with the proteomic analysis, proteasome activity was maximal for both WT and *pfld* leaves at the proliferative stage, and declined thereafter (Fig. 6A). Moreover, differential accumulation of proteasomal subunits correlated with increased activity in *pfld* leaves compared to WT counterparts, but only during the cell expansion phase (Fig. 6A). Involvement of the proteasome in Fld-dependent modulation of leaf size was further supported by the observation that the developmental effects of Fld were suppressed upon exposure of the plants to sub-lethal concentrations of the proteasome inhibitor MG132 (Fig. 6B).

**Fig. 6.**
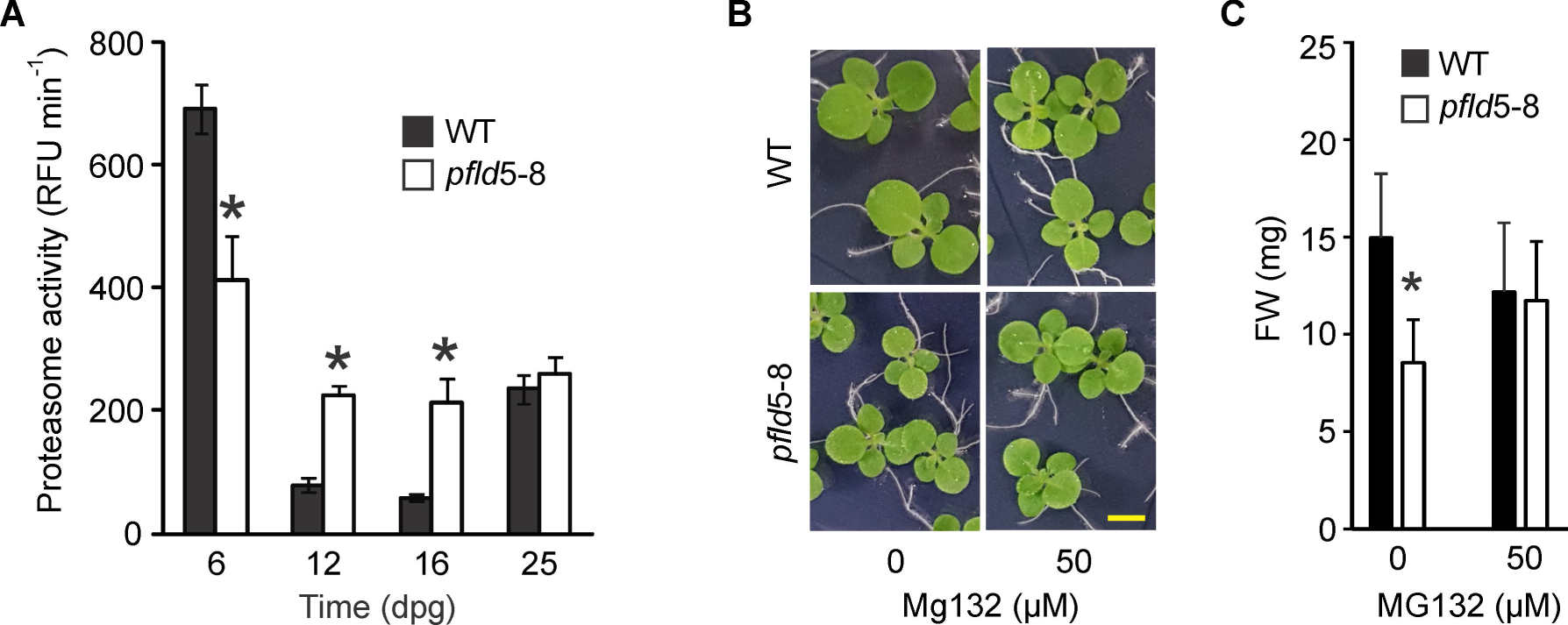
Reversion of the Fld-dependent small size phenotype by treatment with a proteasome inhibitor. (A) Proteasome activity was measured in cleared extracts of leaf 1 at the indicated developmental stages (*n* = 3), using Suc-LLVY-AMC as substrate. RFU, relative fluorescence units. (B) Representative pictures and (C) comparison of shoot growth, measured as fresh weight (FW), in the presence of the proteasome inhibitor MG132 (*n =* 6-8). Bar = 0.5 cm. Data reported are means ± SD. Asterisks indicate statistically significant differences (*P <* 0.05) using one-way ANOVA and Tukey multiple comparison test.

### Decreased cell size in *pfld* leaves was associated with lower endoreduplication

Endoreduplication is a modified version of the cell cycle that lacks the M phase and has been related to increased cell and organ size due to higher ploidy and DNA content in the nuclei (Tsukaya, 2008). We therefore estimated the ploidy levels of fully expanded (25 dpg) leaf 1 of WT and *pfld* plants using flow cytometry. WT nuclei showed a bimodal distribution, with most of its DNA in the 4C form, whereas the majority of *pfld* cells were diploid with lower contribution of 4C (Fig. 7A, B). Accordingly, EI values were significantly lower in *pfld* plants (Fig. 7C), correlating with decreased cell size.

**Fig. 7.**
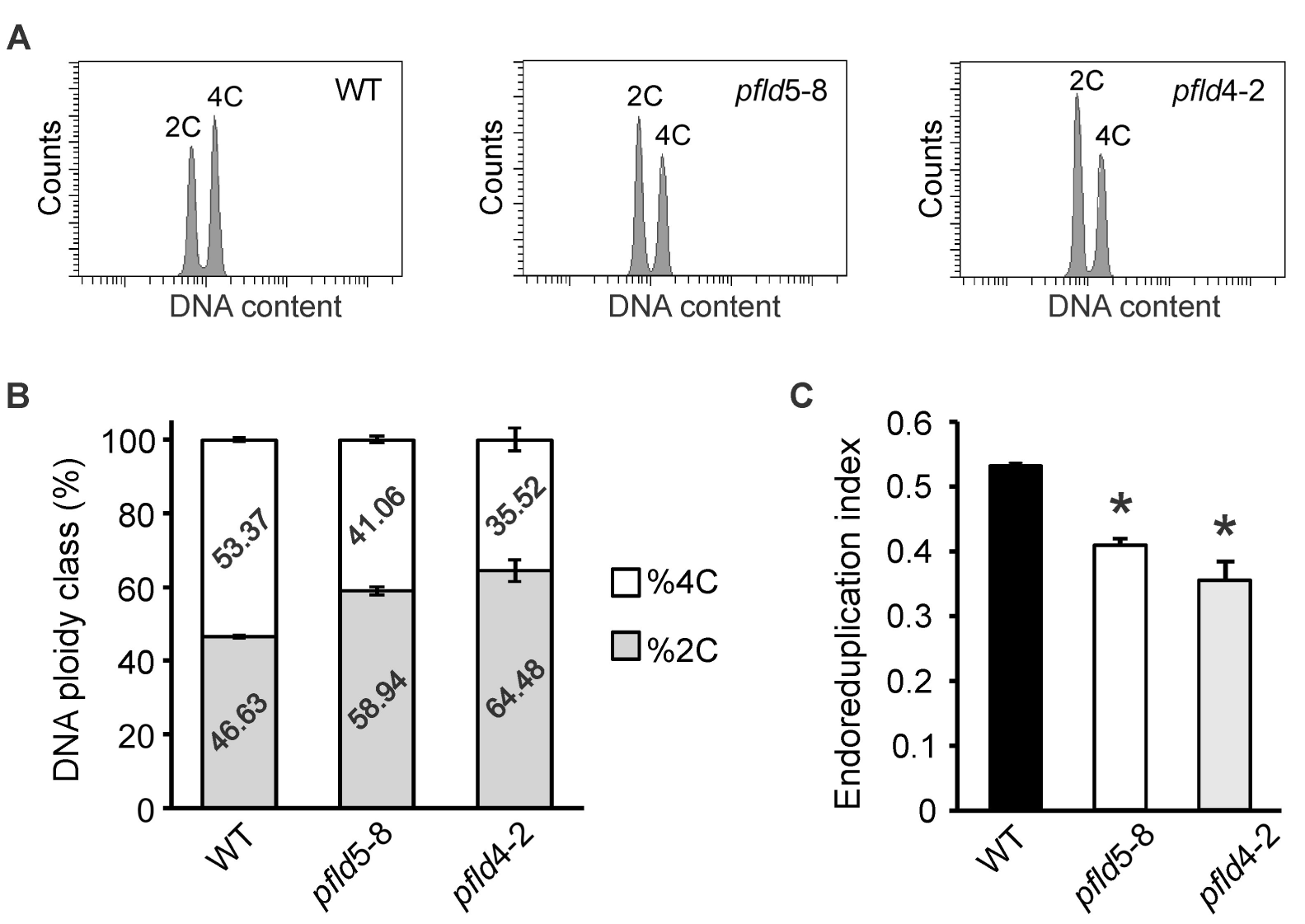
Effect of chloroplast Fld expression on DNA endoreduplication. (A) Flow cytometry profiles of nuclear DNA contents in fully expanded leaf 1 at 25 dpg. (B) Percentage of nuclear DNA contents in the 2C and 4C classes. (C) Endoreduplication index. Data are shown as means ± SD of 3 biological replicates, each consisting of leaf 1 from 10 independent plants per line. Asterisks indicate statistically significant differences with respect to WT plants (*P <* 0.05), as determined using one-way ANOVA and Tukey multiple comparison test.

## Discussion

Leaf development relies on the coordination of cell proliferation and expansion. The first of these processes sets the total number of cells in a leaf, whereas post-mitotic cell expansion determines the final cell size, so that both processes contribute to determine organ size (Gonzalez *et al*., 2012; Kalve *et al*., 2014). The transition from proliferation into expansion proceeds basipetally and occurs quite abruptly during leaf development (Kazama *et al*., 2010; Andriankaja *et al*., 2012). The results obtained using mutants and inhibitors of chloroplast biogenesis indicate that the onset of cell expansion depends on the establishment of a functional photosynthetic machinery and its associated retrograde signal transduction (Andriankaja *et al*., 2012; Hudik *et al*., 2014; van Dingenen *et al*., 2016a). On the other hand, redox-based signals, including ROS generated during photooxidative stress, have been shown to modulate leaf development (Dietz et al., 2016; Foyer and Hanke, 2022). Chloroplasts can make a significant contribution to this process with their rich oxido-reductive metabolism (Juvany *et al*., 2013; Muñoz and Munné-Bosch, 2018). We show here that introduction of the alternative electron sink Fld, which is known to effectively relieve the excitation pressure on the PETC (Gómez *et al*., 2020), resulted in decreased leaf size in tobacco plants (Fig. 1; Supplementary Fig. S1). We therefore propose that when present in chloroplasts, Fld interferes with retrograde redox signaling involved in leaf development.

The oxidative burst that occurs during early leaf development has been associated to defective PETC function, but the mechanisms responsible for this transient ROS accumulation are poorly understood. Proteomic analysis showed that light-harvesting proteins began to accumulate before electron-transfer components (Fig. 3; Supplementary Fig. S10, S11). Enzymes associated with *Chl* metabolism also began to increase their levels from the early stages of leaf development (Supplementary Fig. S12), in line with previous reports (Andriankaja *et al*., 2012; Omidbakhshfard *et al*., 2021). The sequential nature of the PETC assembly provides a plausible explanation for the oxidative explosion. If light harvesting is put into operation while the photosynthetic machinery is still under construction and the energy dissipation mechanisms are not yet fully functional, it is likely that a significant proportion of the absorbed photons cannot be used for productive photochemistry and instead participate in energy and electron transfer to oxygen. This in turn leads to increased ROS propagation in WT immature plastids, as shown in Fig. 4 and in the schematic model of Fig. 8A. The scheme illustrates the negative correlation between photosynthetic activity and ROS build-up (Juvany *et al*., 2013; Muñoz and Munné-Bosch, 2018). By accepting reducing equivalents from the PETC, Fld is proposed to limit ROS accumulation (Fig. 4C, 8A) and to increase the oxidation state of key signaling components of the PETC such as the plastoquinone pool (Gómez *et al*., 2020). It is worth noting, within this context, that Fld does not simply act as a dissipative electron sink device such as the Mehler-Asada cycle or chlororespiration, but instead it delivers the surplus of reducing equivalents to productive redox pathways of the chloroplast.

**Fig. 8.**
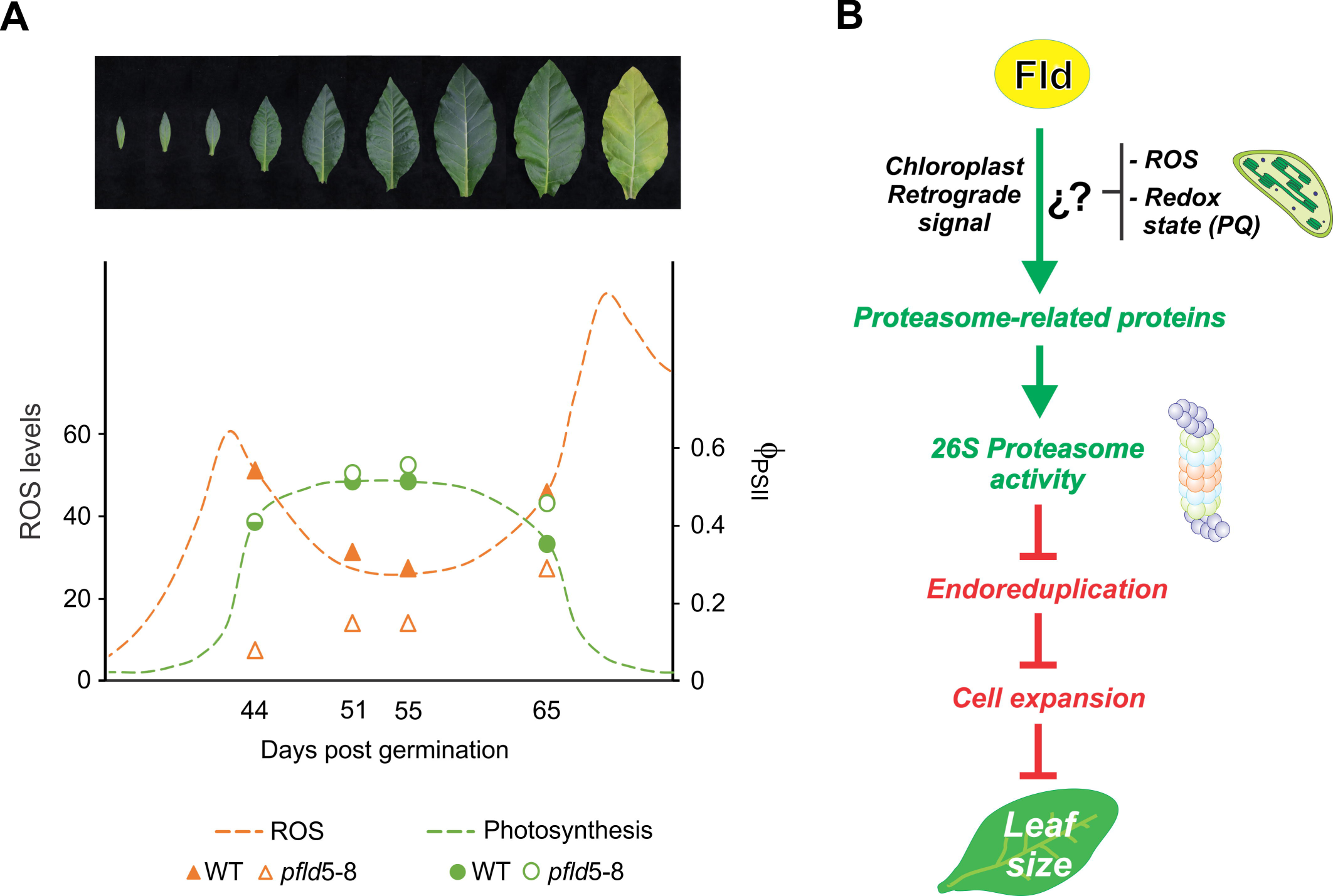
Proposed model for Fld action in leaf development. (A) Schematic representation of photosynthesis (green) and ROS production (orange) during leaf development. Dotted lines were drawn based on similar models reported by Juvany *et al*. (2013) and Muñoz and Munné-Bosch (2018). Experimental data points for ROS levels (triangles) and __PSII_ (circles) were taken from Fig. 4 and overimposed for comparison. Filled and empty symbols represent WT and pfld leaves, respectively. (B) Proposed model for the effect of Fld on leaf development. Fld activity as electron sink modifies chloroplast redox poise (Gómez *et al*., 2020) and ROS production (Fig. 4C), altering retrograde signalling output. A consequence of this modification is induction of proteasome expression (Fig. 5) and activity (Fig. 6A). It is likely that the effects of increased proteasome activity on leaf size involve inhibition of DNA endoreduplication, as reported previously (Sako *et al*., 2010).

Since leaf size can be modified by the addition of sugars, the role of photosynthesis has been often associated with the provision of carbohydrates, not only as a source of nutrients and energy but also as signals (Smeekens and Hellmann, 2014; van Dingenen *et al*., 2016b). Thus, the Fld effect could be mediated by enhanced carbon metabolism resulting from incorporation of the additional electron carrier into the photosynthetic process. Metabolic profiling of mature *pfld* leaves from plants grown under normal conditions suggests otherwise: steady-state levels of soluble carbohydrates such as sucrose, glucose and fructose, were not affected by the activity of the flavoprotein in chloroplasts (Tognetti *et al*., 2007: Mayta *et al*., 2018). Moreover, Fld and sugars affect organ development in different ways. Sucrose, for instance, increases final leaf size by promoting cell proliferation and postponing exit from this phase (van Dingenen *et al*., 2016b), whereas Fld specifically decreases the rates of cell expansion, without affecting the timespan of the phases (Fig. 2; Supplementary Fig. S4). Also, plant exposure to sugars resulted in smaller chloroplasts in mesophyll cells (van Dingenen *et al*., 2016b). Fld expression, instead, had no significant effect on plastid size or ultrastructure (Supplementary Fig. S9). Association of Fld activity to redox metabolism and ROS control (rather than carbohydrate synthesis) also concurs with the observation that antisense Arabidopsis plants deficient in different peroxiredoxin isoforms (lower antioxidant capacity) exhibit enlarged leaves and a giant phenotype (Jeong *et al*., 2022). Incidentally, they also display accelerated leaf senescence (Jeong *et al*., 2022). Although further research is necessary to properly settle this issue, the results suggest that Fld impact on leaf development was not associated to sugar signaling.

As a first step to unravel the mechanisms by which chloroplast redox signals might control leaf cell expansion it is necessary to identify their cellular targets. Previous observations using transcriptional profiling of *pfld* plants grown under normal conditions revealed that levels of over 1,000 leaf transcripts were differentially affected by the presence of the flavoprotein, with the generalized build-up of RNAs encoding proteasomal subunits being the most remarkable feature of the plant response to Fld expression (Pierella-Karlusich *et al*., 2017, 2020). This transcriptional up-regulation (Supplementary Figure S14) was reflected in the protein accumulation profiles (Fig. 5), which also showed that the differential induction of proteasomal components in response to Fld presence started at the expansion phase of growth. Levels of proteasomal proteins in WT leaves were maximal during proliferation and declined thereafter (Supplementary Fig. S15). This result is supported by the observation that proteasome activity was highest at this developmental stage (Fig. 6A). The early induction of proteasomal subunits and activity during leaf development has been reported previously (Andriankaja *et al*., 2012; Omidbakhshfard *et al*., 2021), and attributed to the need of coping with the surplus of unfolded targets resulting from high rates of protein synthesis during active cell and organelle division. Indeed, protein translation was identified as the main category enriched in proliferating Arabidopsis cells (Omidbakhshfard *et al*., 2021).

When present, chloroplast Fld partially prevented the age-dependent decline of proteasomal components, leading to their differential accumulation relative to WT plants (Fig. 5; Supplementary Fig. S14). Interestingly, the 20S core particle responded very strongly to the presence of Fld, compared to the 19S regulatory complex or other ancillary components of the 26S proteasome (Fig. 5; Supplementary Fig. S15). This differential build-up correlated with the concomitant increase of proteasome activity in *pfld* leaf extracts, which also initiated upon transition into the expansion phase (Fig. 6A), namely, the same developmental stage at which chloroplast Fld exerted its differential effect on leaf growth (Fig. 2A; Supplementary Fig. S4).

Several lines of evidence support the central role played by protein degradation via the proteasome in the determination of organ size. Pharmacological approaches and analysis of loss-of-function mutants indicate that higher proteasome activity decreases leaf size, and *vice versa* (Kurepa and Smalle, 2008; Sonoda *et al*., 2009; Nguyen *et al*., 2013). Interestingly, different components of the 26S complex were shown to operate at various levels of the organ development pathway. For instance, plants deficient in the ubiquitin receptor DA1 or the E3 ligase BIG BROTHER, exhibit leaf enlargement caused by changes in the rate and duration of the proliferative phase (Disch *et al*., 2006; Vanhaeren *et al*., 2017), whereas inactivation of the AAA ATPase RPT2a, belonging to the 19S regulatory subunit, led to a similar outcome by increasing cell expansion through extended endoreduplication (Kurepa *et al*., 2009; Sonoda *et al*., 2009).

A global induction of proteasomal components similar to that elicited by Fld has been reported upon constitutive expression of the upstream positive regulator RPX, which also resulted in smaller leaves (Nguyen *et al*., 2013). Involvement of proteasome function in the phenotypes elicited by Fld and RPX was confirmed by partial reversion of the leaf size phenotype using specific inhibitors of proteasome activity (Fig. 6B; Nguyen *et al*., 2013). Leaf growth reduction in *pfld* tobacco plants was accompanied by a decline in DNA polyteny (Fig. 7), a correlation already reported for Arabidopsis mutants deficient in proteasome activity (Sonoda *et al*., 2009; Sako and Yamaguchi, 2010).

As a working hypothesis, we thus propose the sequence of events depicted in the model of Fig. 8B: redox-based signals transmitted by chloroplasts influence proteasome expression and activity, which in turn modulate cellular processes involved in organ size determination, including endoreduplication (Fig. 8B). Redox regulation of the proteasome is usually associated with oxidative rather than reductive stimuli (Lefaki *et al*., 2017), although NADH has been shown to stabilize the 26S complex in animal cells (Tsvetkov *et al*., 2014). Redox signals generated in chloroplasts can be both oxidative (i.e., ROS) or reductive (namely, over-reduction of the PETC), and Fld could mitigate both of them. Further studies will be required to distinguish which of these possible signals is affecting proteasome expression and function during leaf development. For instance, a combination of alternative electron sinks accepting electrons from different components of the PETC might help to dissect the nature of the redox signaling relay.

In summary, we report here that tailored changes in chloroplast redox status have a direct impact on final leaf size (Fig. 8). Research on chloroplast retrograde regulation has gained momentum in recent years, and has led to the identification of several plastid-generated signals that can influence nuclear gene expression (He *et al*., 2018). These developmental cues likely interact in a cooperative way to modulate organ development. The use of alternative electron transport shuttles, as illustrated here, provides novel tools to investigate how this multiplicity of signaling pathways coordinate and cross-talk in the determination of leaf size and shape, which constitutes a most exciting goal of plant biology with potential impact in agriculture.

### Supplementary data

The following supplementary data are available at JXB online.

**Fig. S1.** Phenotypes of WT and Fld-expressing plants along development.

**Fig. S2.** Effect of a plastid-targeted Fld on leaf epidermal cell size, cell numbers and stomatal index.

**Fig. S3.** Influence of plastid Fld on the size and number of proliferating cells.

**Fig. S4.** Effect of chloroplast-targeted Fld on final size of leaf 10.

**Fig. S5.** Variations in cell density during leaf development in WT and Fld-expressing plants.

**Fig. S6.** Changes in photosynthetic pigments per area during development of WT and *pfld* leaves.

**Fig. S7.** Changes in photosynthetic pigments per cell during development of WT and *pfld* leaves.

**Fig. S8.** Effect of Fld expression on plastid chlorophyll contents and chloroplast numbers per cell.

**Fig. S9.** Ultrastructural features of chloroplasts from WT and *pfld* leaves.

**Fig. S10.** Expression of photosynthesis-related proteins through development in tobacco *pfld*5-8 plants.

**Fig. S11.** Expression of photosynthesis-related proteins through development in tobacco *pfld*4-2 plants.

**Fig. S12.** Expression of proteins related to chlorophyll synthesis during development of WT and *pfld* leaves.

**Fig. S13.** Light-response curves of photosynthesis-associated parameters in developing WT and *pfld* leaves.

**Fig. S14.** Expression of proteasomal components during development of WT and *pfld* leaves.

**Fig. S15.** Effect of a plastid-targeted Fld on the expression of the proteasome system along leaf development.

**Table S1.** Primers used for quantitative real-time PCR determinations.

## Acknowledgements

The authors wish to thank Diego Aguirre for his technical assistance with plant growth at IBR. We also acknowledge the service of the Mass Spectrometry Unit of the Institute of Molecular and Cellular Biology of Rosario (UEM-IBR) to perform the proteomic experiment, and are indebted to Dr. Eduardo Ceccarelli, Dr. Germán Rosano and Alejo Cantoia of UEM-IBR for the many contributions to the analysis and interpretation of the retrieved data. We thank Marion Benecke and Kirsten Hoffie (IPK Gatersleben) for technical assistance with sample preparation for histology and electron microscopy. We also acknowledge Dr. Raquel Chan (IAL Santa Fe, Argentina) for the use of the LI-6400 portable photosynthesis system

## Author contributions

NC, AFL and MLM: conceptualization; RCA, MLM, MM and M-RH: methodology: RCA, MLM, MM, M-RH, AFL and NC: investigation and formal analysis; RCA, MLM, MM, M-RH, AFL and NC: writing - review & editing; MM, M-RH, AFL and NC: resources. All authors have read and agreed to the published version of the manuscript.

## Data availability

The proteomics data underlying this article are available in ProteomeXchange, and can be accessed with with identifier PXD041855.

## Abbreviations

AMC: 7-amino-4-methylcoumarin
DEPs: differentially expressed proteins
dpg: days post germination
FC: fold-change
Fd: ferredoxin
Fld: flavodoxin
KEGG: Kyoto Encyclopedia of Genes and Genomes
PETC: photosynthetic electron transport chain
φ_PSI_: quantum yield of PSI
φ_PSII_: quantum yield of PSII
ROS: reactive oxygen species
WT: wild-type

