## Supplementary data for "Introduction of a terminal electron sink in chloroplasts decreases leaf cell expansion associated to higher proteasome activity and lower endoreduplication"

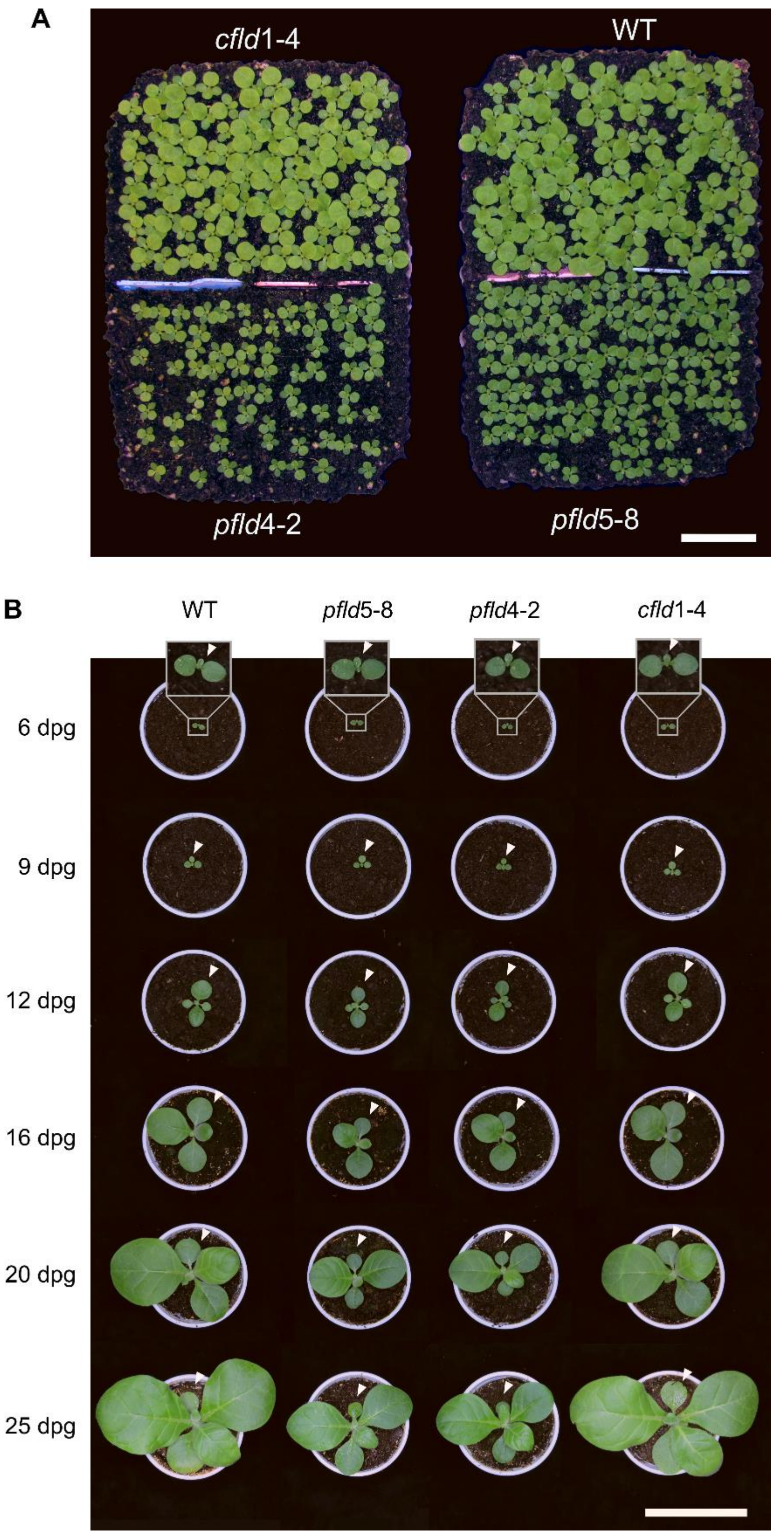


**Supplementary Fig. S1. Development of WT and Fld-expressing plants.** Plants were grown under chamber conditions (Materials and methods). (A) Phenotypes of WT, *pfld* and *cfld* plants at 14 dpg. Bar = 5 cm. (B) Pictures of the various lines were taken at the indicated days post germination (dpg). Arrowheads show leaf 1. Bar = 10 cm.

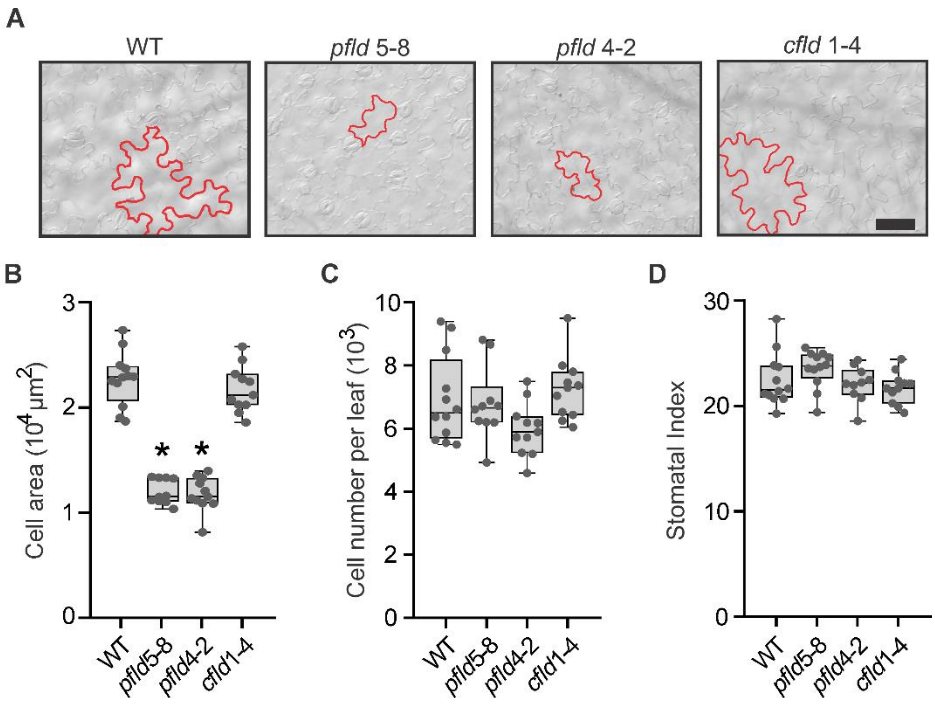


**Supplementary Fig. S2. Effect of plastid-targeted Fld on leaf epidermal cell size, cell numbers and stomatal index.** (A) Representative micrographs from epidermal cells of leaf 1 from WT, *pfld* and *cfld* tobacco plants at 35 dpg. Contours of typical cells are shown in red. Bar = 100 µm. (B) Cell areas, (C) cell numbers per leaf, and (D) stomatal index were determined from clarified leaf tissue as described in Materials and methods. Four regions of the leaf blade (Figure 1D) were used, and their results averaged. Data are presented as box plots ($n$ = 10-12 biological replicates per genotype). Central horizontal lines show the medians; box limits indicate the 25$^{th}$ and 75$^{th}$ percentiles; whiskers extend 1.5-times the interquartile range from the 25$^{th}$ and 75$^{th}$ percentiles; closed circles represent data points. Asterisks indicate statistically significant differences with respect to WT siblings ($P < 0.05$), as determined using one-way ANOVA and Tukey multiple comparison test or Kruskall–Wallis one-way ANOVA and Dunn's multiple range test.

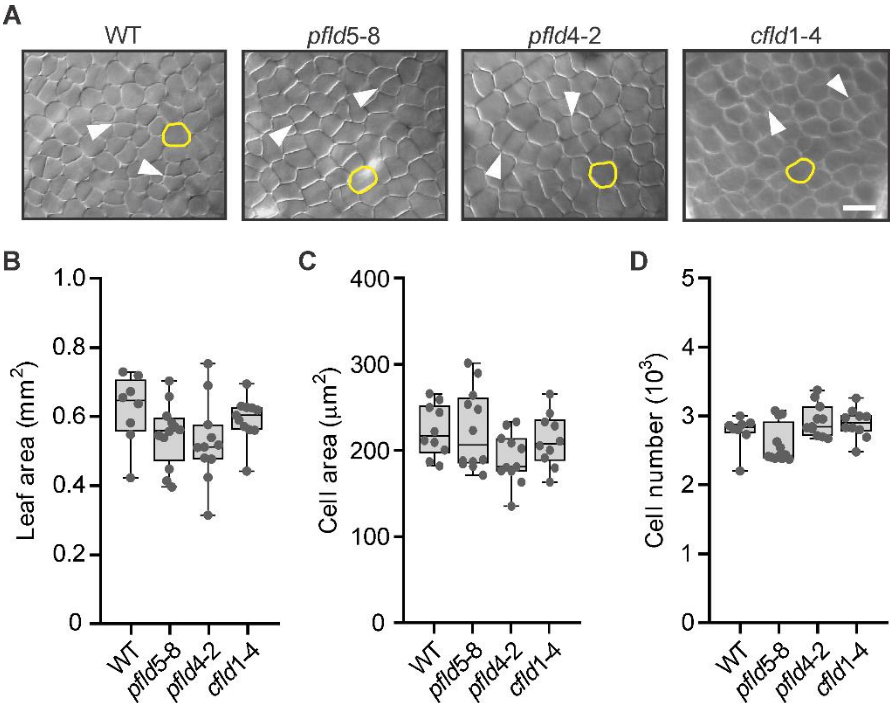


**Supplementary Fig. S3. Influence of plastid Fld on the size and number of proliferating cells.** (A) Representative micrographs from palisade parenchyma cells of leaf 1 from WT, *pfld* and *cfld* tobacco plants at 6 dpg. White arrows indicate cells in division. Contour of typical cells are shown in yellow. Bar = 25 µm. (B) Leaf 1 areas, (C) cell areas and (C) cell numbers per leaf were determined as described in Materials and methods. Four regions of the leaf blade (Figure 1D) were used, and their measurements averaged. Data are reported as box plots ($n$ = 8-12 biological replicates per genotype). Central horizontal lines show the medians; box limits indicate the 25$^{th}$ and 75$^{th}$ percentiles; whiskers extend 1.5-times the interquartile range from the 25$^{th}$ and 75$^{th}$ percentiles; closed circles represent data points. Asterisks indicate statistically significant differences with respect to WT siblings ($P$ < 0.05), determined using one-way ANOVA and Tukey multiple comparison test.

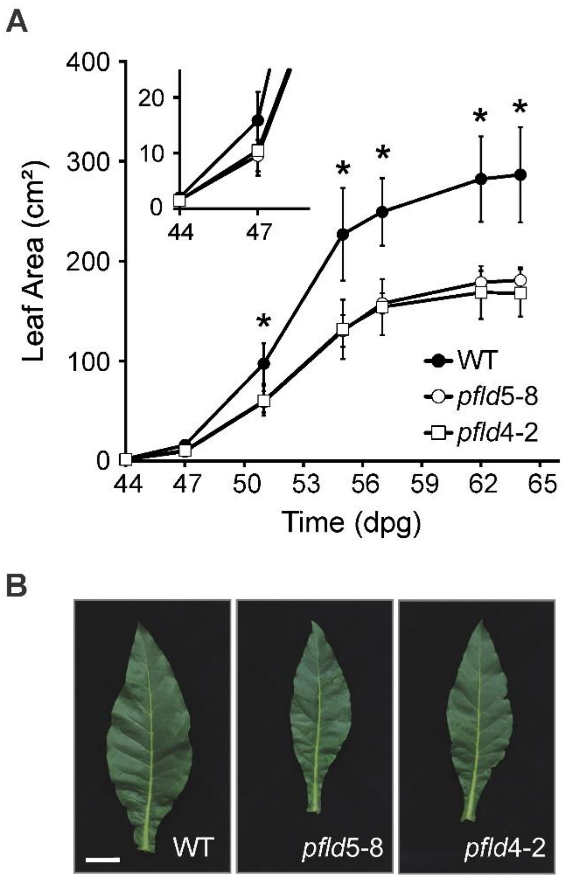


**Supplementary Fig. S4. Effect of chloroplast-targeted Fld on leaf 10 size.** (A) The areas of leaf 10 from WT and *pfld* plants were determined at the indicated dpg. Areas are reported as means ± SD of 11-15 independent plants per line. Asterisks indicate statistically significant differences between WT and both *pfld* lines ($P < 0.05$), determined using one-way ANOVA and Tukey multiple comparison test. (B) Phenotypes of typical leaf 10 from WT and *pfld* plants at the end of the experiment (64 dpg). Bar = 5 cm.

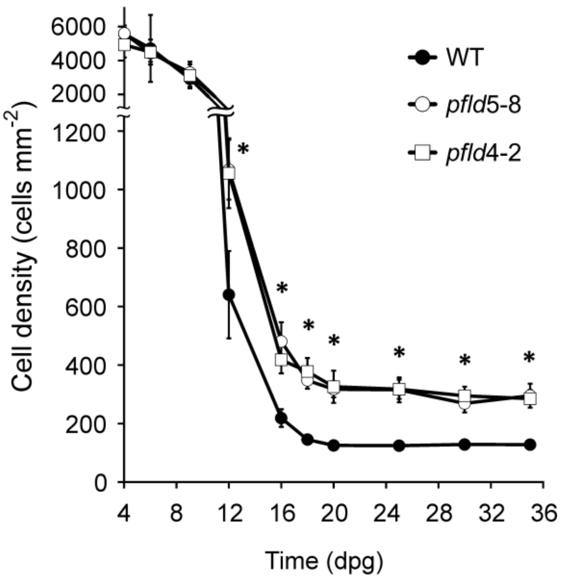


**Supplementary Fig. S5. Variations in cell density during leaf development in WT and Fld-expressing plants.** Cell density was calculated as the ratio of cell numbers over leaf area for leaf 1 of WT and *pfld* tobacco plants at the indicated dpg. Data are reported as means ± SD from at least 8 biological replicates. Asterisks indicate statistically significant differences between WT and both *pfld* lines ($P < 0.05$), determined using one-way ANOVA and Tukey multiple comparison test.

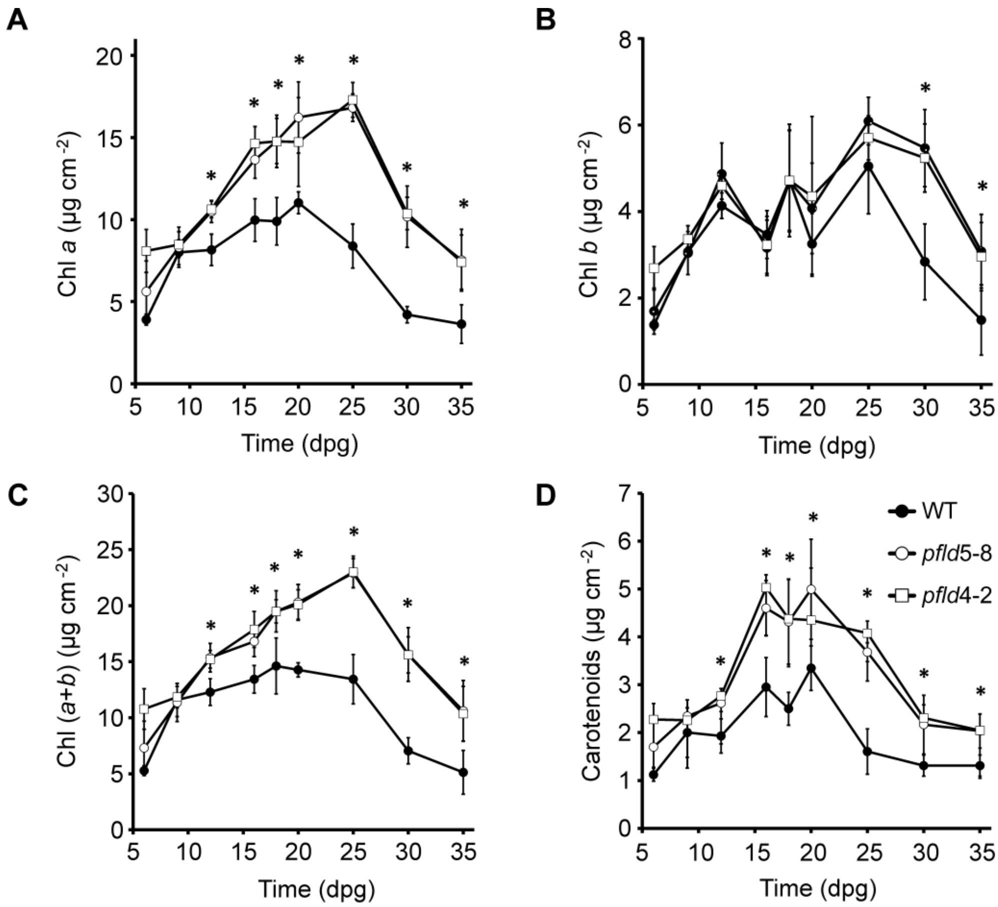


**Supplementary Fig. S6. Changes in photosynthetic pigments per area during development of WT and *pfld* leaves.** Contents of (A) *Chl a*, (B) *Chl b*, (C) *Chl (a+b)*, and (D) carotenoids were determined in leaf 1 of WT and *pfld* plants at the indicated dpg**.** Data are shown as means ± SD of at least 5 biological replicates, each one comprising at least 8 independent leaves per genotype. Asterisks indicate statistically significant differences between WT and both *pfld* lines ($P < 0.05$), determined using one-way ANOVA and Tukey multiple comparison test.

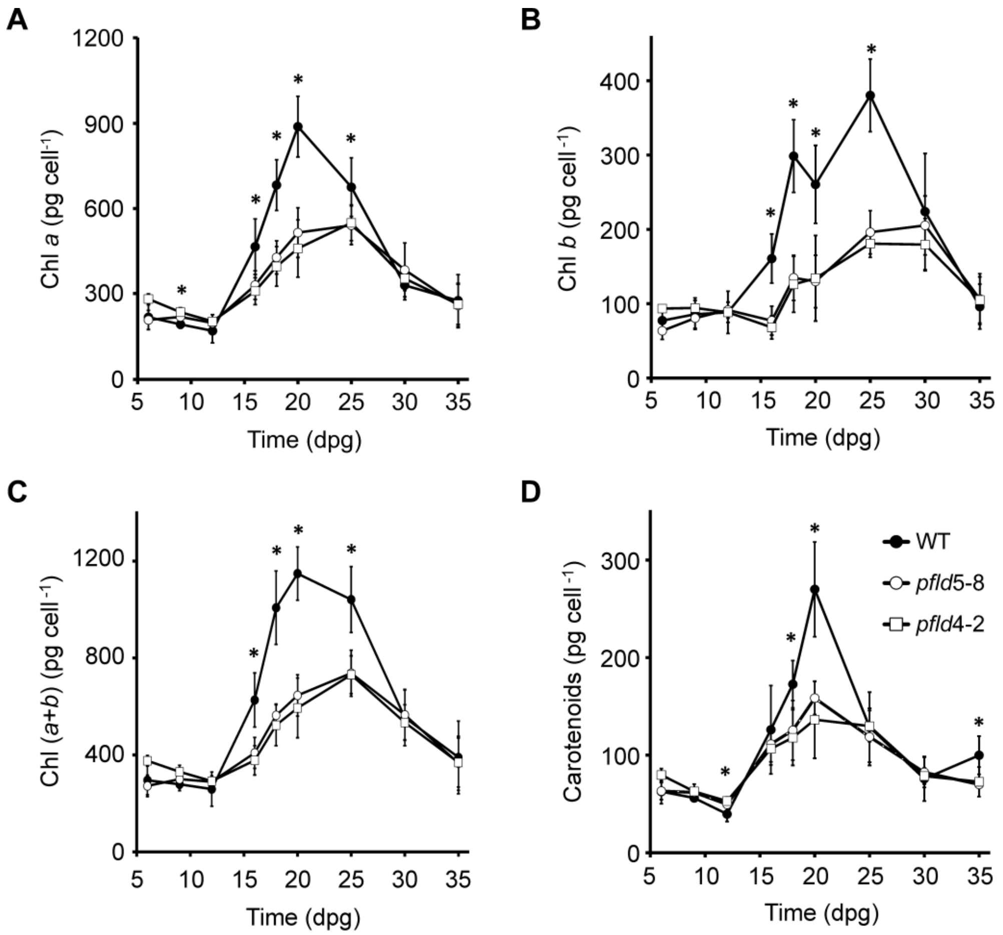


**Supplementary Fig. S7. Changes in photosynthetic pigments per cell during development of WT and *pfld* leaves.** Contents of (A) *Chl a*, (B) *Chl b*, (C) *Chl (a+b)*, and (D) carotenoids per leaf area (Figure S6) were divided by the cell numbers per leaf area (Figure 2C) to obtain the corresponding values of pigment contents per cell. All other conditions are those of Supplementary Figure S6.

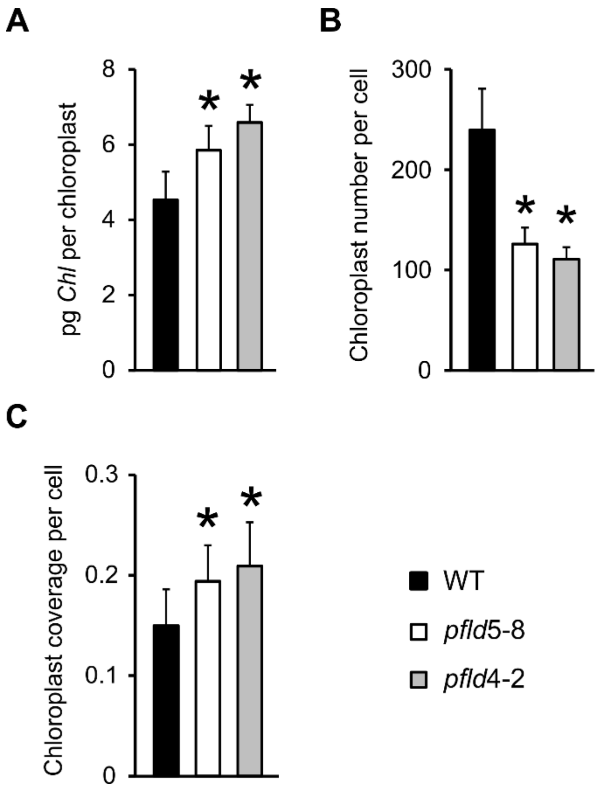


**Supplementary Fig. S8. Effect of Fld expression on plastid chlorophyll contents and chloroplast numbers per cell.** (A) Total *Chl* content per chloroplast in mature leaf 1 from WT and *pfld* plants at 25 dpg. Chloroplasts were isolated and counted as described in Materials and methods. Data are means ± SD of 7-8 independent plants per line. (B) Chloroplast numbers per cell were estimated from *Chl* (*a*+*b*) per cell (Supplementary Figure S7C) and total *Chl* per chloroplast (A). Data are means ± SD of 7-8 independent plants per line. (C) Chloroplast coverage was calculated as the ratio between total chloroplast area and cell area (Figure 2B) as described in Materials and methods. Data reported are means ± SD of 47-52 biological replicates per genotype. Asterisks indicate statistically significant differences ($P$ < 0.05), as determined using one-way ANOVA and Duncan's multiple comparison test.

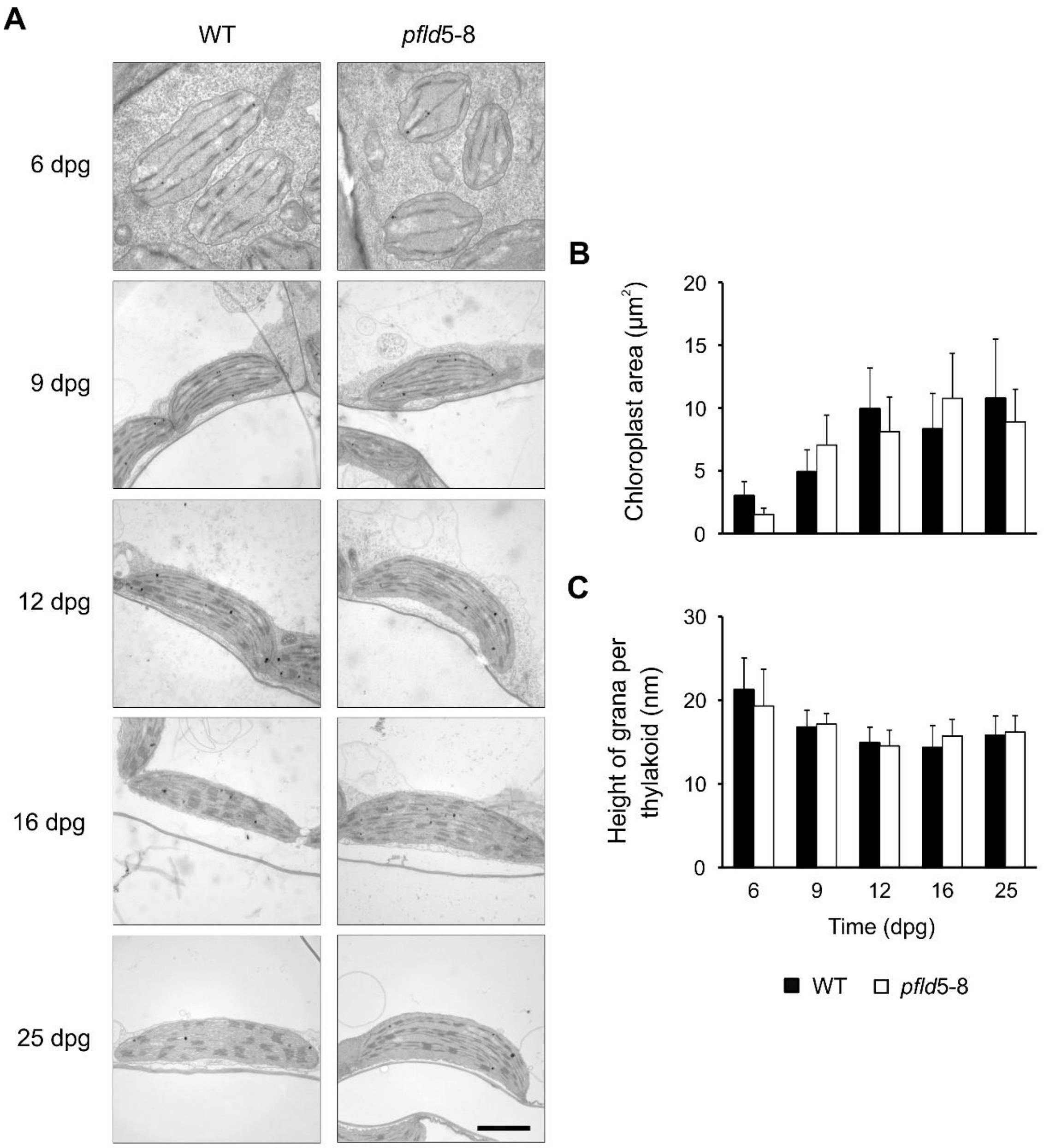


**Supplementary Fig. S9. Ultrastructural features of chloroplasts from WT and *pfld* leaves.** (A) Representative electron micrographs of palisade parenchyma tissue showing typical chloroplasts from leaf 1 of WT and *pfld*5-8 plants at the indicated dpg. Bar = 500 nm. (B) Average chloroplast area, and (C) grana height as determined from the transmission electron microscopic images by image analysis (see Materials and methods). Data are shown as means ± SD of 50-100 biological replicates.

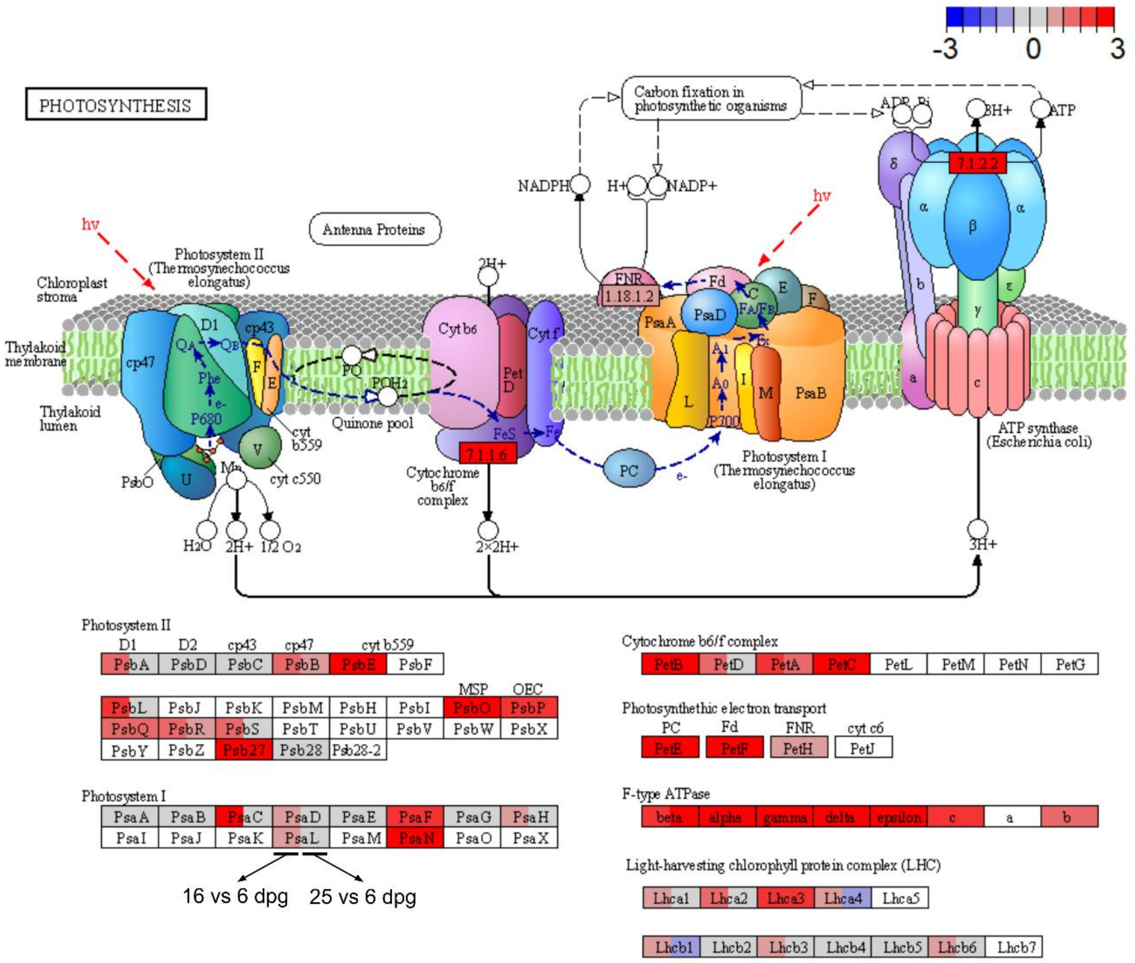


**Supplementary Fig. S10. Expression of photosynthesis-related proteins through development in tobacco *pfld*5-8 plants.** Analysis of KEGG pathway changes was carried out using Pathview. The color scale indicates the extent of up- (red) or down-regulation (blue). The upper part of the figure shows a scheme of the PETC with its multiple components (each displaying the acronym of its name), whose regulation is detailed in the lower part. Rectangles in the upper panel indicate the activities of selected complexes and enzymes as defined by the IUBMB: EC 7.1.1.6, plastoquinone-plastocyanin reductase; EC 1.18.1.2, ferredoxin-NADP$^+$ reductase; EC 7.1.1.2, ATP synthase. Rectangles in the lower panel show the modulation of individual subunits. Each rectangle (upper and lower panels) was divided into 2 sections, which represent proteins modulated at 16 dpg vs 6 dpg (left), and 25 dpg vs 6 dpg (right). Empty boxes correspond to components that were not detected in the proteomic analysis. Activities depicted in the upper panel are suggested to be induced at the two developmental stages according to Pathview, as indicated by the red filling. Details on the experimental design and data analysis are given in Materials and methods.

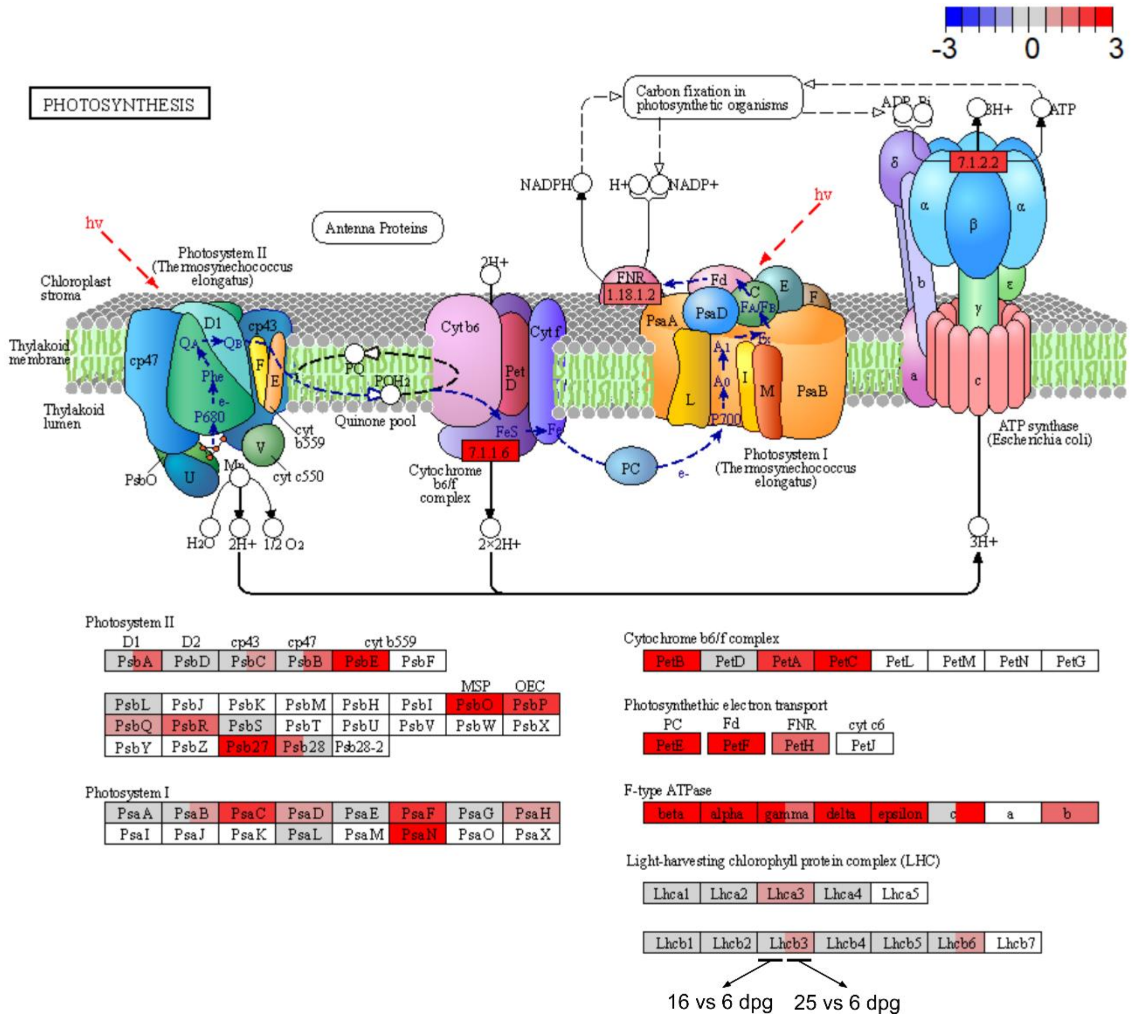


**Supplementary Fig. S11. Expression of photosynthesis-related proteins through development in tobacco *pfld*4-2 plants.** Analysis of KEGG pathway changes was carried out using Pathview. The color scale indicates the extent of up- (red) or down-regulation (blue). The upper part of the figure shows a scheme of the PETC with its multiple components (each displaying the acronym of its name), whose regulation is detailed in the lower part. Rectangles in the upper panel indicate the activities of selected complexes and enzymes as defined by the IUBMB: EC 7.1.1.6, plastoquinone-plastocyanin reductase; EC 1.18.1.2, ferredoxin-NADP$^+$ reductase; EC 7.1.1.2, ATP synthase. Rectangles in the lower panel show the modulation of individual subunits. Each rectangle (upper and lower panels) was divided into 2 sections, which represent proteins modulated at 16 dpg vs 6 dpg (left), and 25 dpg vs 6 dpg (right). Empty boxes correspond to components that were not detected in the proteomic analysis. Activities depicted in the upper panel are suggested to be induced at the two developmental stages according to Pathview, as indicated by the red filling. Details on the experimental design and data analysis are given in Materials and methods.

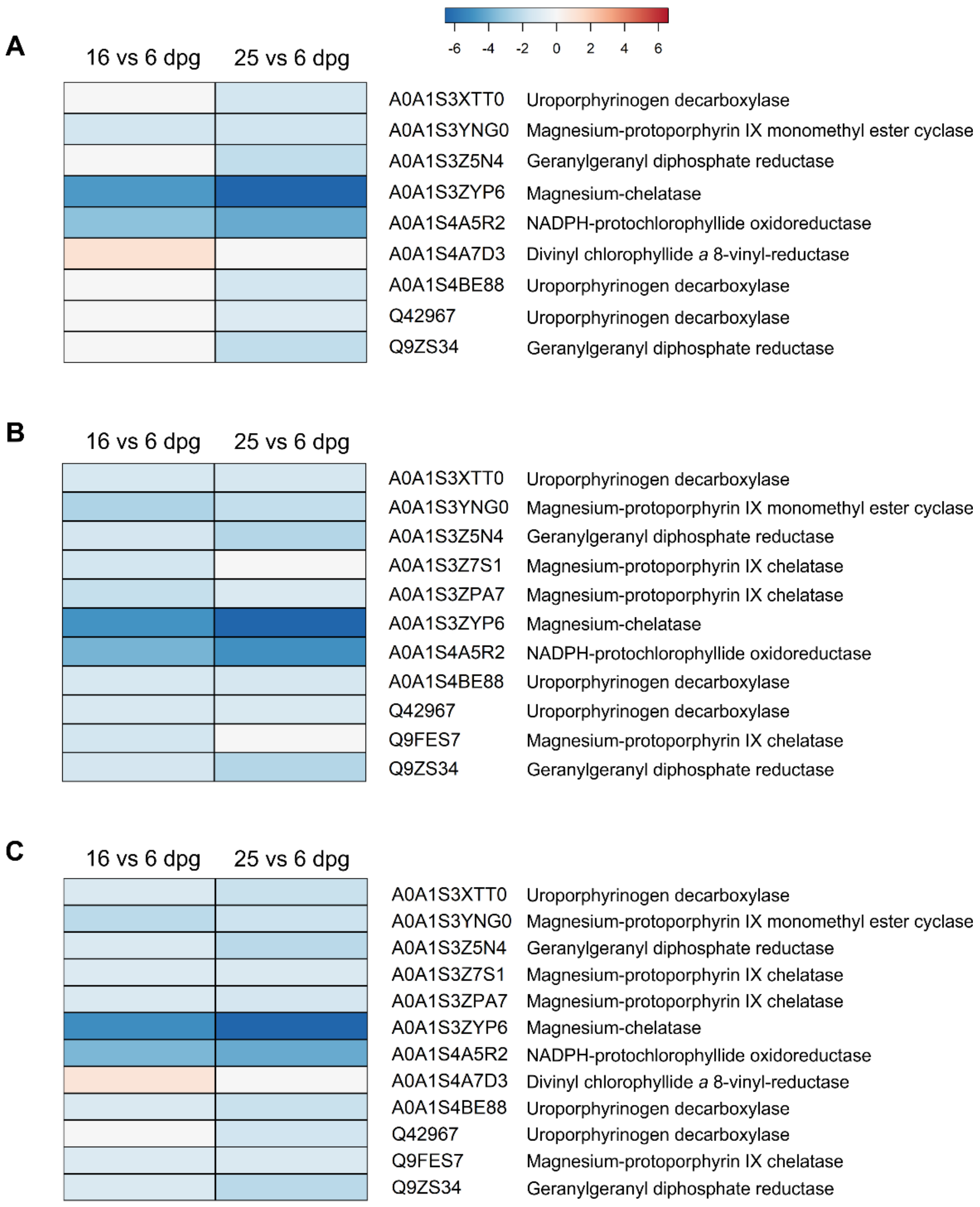


**Supplementary Fig. S12. Expression of proteins related to chlorophyll synthesis during development of WT and *pfld* leaves.** Heat map of DEPs abundance at 16 dpg vs 6 dpg and 25 dpg vs 6 dpg for WT (A), *pfld*5-8 (B) and *pfld*4-2 (C) leaves. The list of proteins used for this analysis belongs to the biological process "chlorophyll biosynthesis" from Unitprot (ID KW-0149). The color scale indicates the extent of up- (red) or down-regulation (blue). Details on the proteomics assay and analysis are given in Materials and methods.

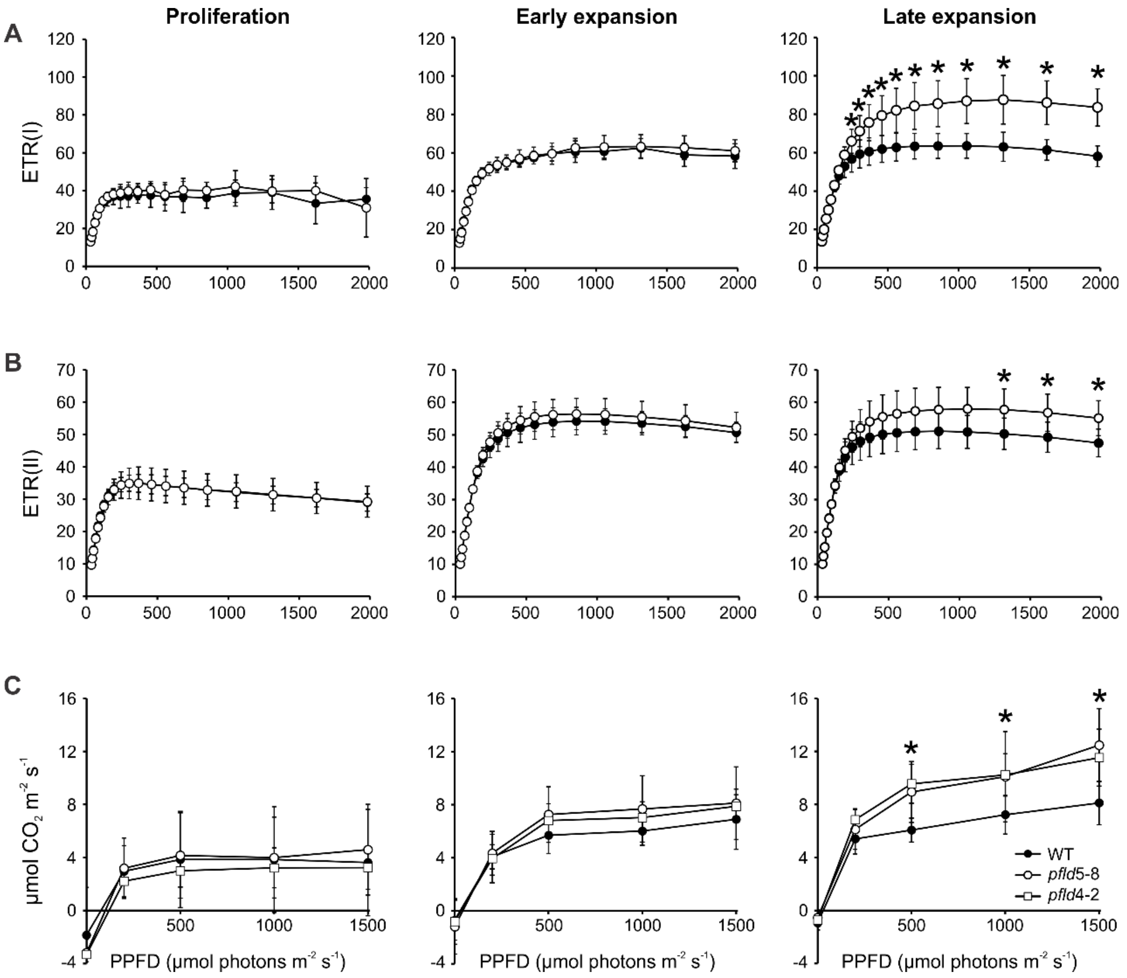


**Supplementary Fig. S13. Light-response curves of photosynthesis-associated parameters in developing WT and *pfld* leaves.** Determinations of (A) light-dependent electron transport rates of PSI, (ETR(I)), (B) light-dependent electron transport rates of PSII (ETR(II)), and (C) net $CO_2$ uptake rates, as a function of photosynthetic photon flux density (PPFD), were carried out in leaf 10 at proliferation (44 dpg), early expansion (51 dpg) and late expansion (55 dpg) phases ($n$ = 5-7). Data reported are means ± SD, and asterisks indicate statistically significant differences ($P < 0.05$), determined using one-way ANOVA and Duncan's multiple comparison test.

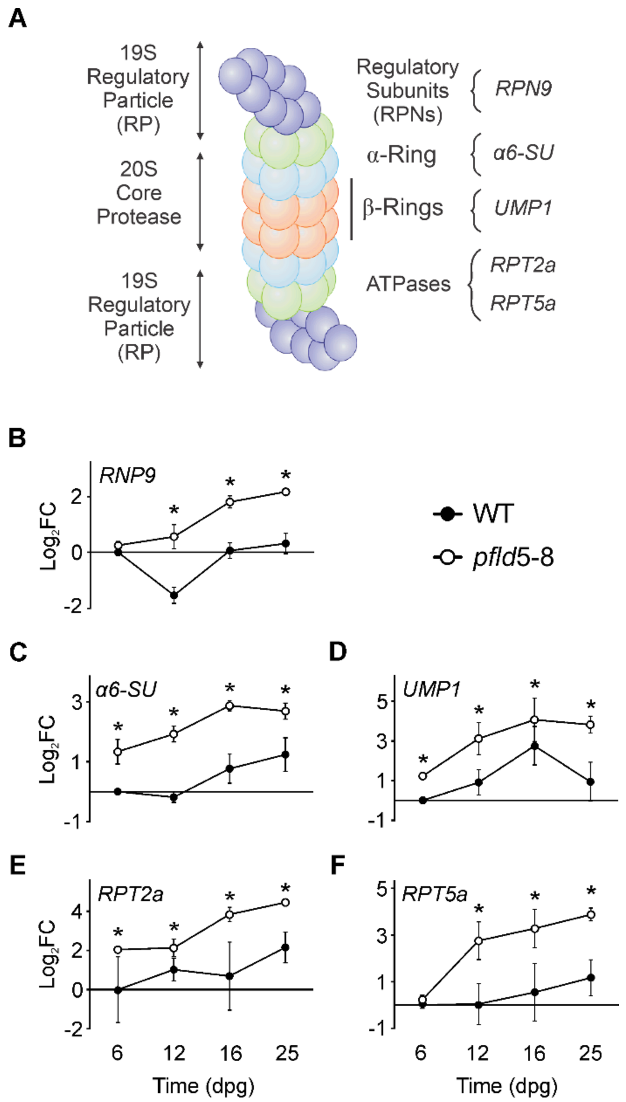


**Supplementary Fig. S14. Induction of proteasome-encoding genes during development of WT and *pfld* leaves.** (A) Scheme of the plant proteasome structure showing selected target genes with its reported location. Transcript levels of the (B) *N9 subunit of the 19S Regulatory Particle of the proteasome* (*RPN9*), (C) *Alpha 6 subunit of the 20S Core Protease of the proteasome* (*α6-SU*), (D) *Proteasome maturation protein 1* (*UMP1*), (E) *Regulatory Particle AAA-ATPase 2a* (*RPT2a*), and (F) *Regulatory Particle AAA-ATPase5a* (*RPT5a*) were determined in leaf 1 of WT and *pfld*5-8 plants at the indicated dpg by qRT-PCR. Values are means ± SD of fold-changes in $\log_2$ scale relative to the levels estimated in leaf 1 from WT plants at 6 dpg ($n$ = 6). Asterisks indicate statistically significant differences ($P$ < 0.05), as determined using one-way ANOVA and Tukey multiple comparison test.

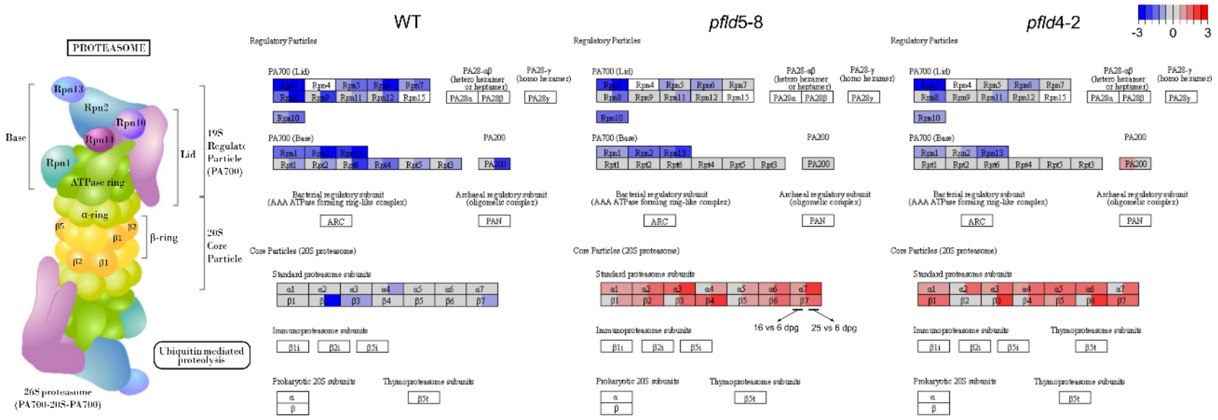


**Supplementary Fig. S15. Effect of plastid-targeted Fld on the down-regulation of the proteasome system along leaf development.** Analysis of KEGG pathway changes was carried out using Pathview. Each rectangle was divided into two subsections, which represent proteins modulated at 16 dpg vs 6 dpg and 25 dpg vs 6 dpg, respectively. The color scale indicates the extent of up- (red) or down-regulation (blue). Empty boxes indicate components not detected in the proteomic analysis. Further experimental details are given in Materials and methods.

**Table S1.** Primers used for qRT-PCR determinations.

| Gene | Abbreviation | Accession number | Sequence (5'-3') | Tm (°C) | Product length (bp) | Reference |
|---|---|---|---|---|---|---|
| *N.tabacum* RPN9 subunit of the 19S Regulatory Particle of the proteasome | *RPN9* | Nta.5649 | ACCATAAAGCTCGCCAGGAGTT | 59 | 142 | Pierella Karlusich *et al.*, 2017 |
| | | | TTGTCTCCCAGCAAAGCTGACA | 60 | | |
| *N.tabacum* Proteasome maturation protein 1 | *UMP1* | Nta.2154 | CGGGCGCAATACCATCTTCAAT | 59 | 126 | Pierella Karlusich *et al.*, 2017 |
| | | | GATGCGTGTCAACTGGACGAAA | 59 | | |
| *N.tabacum* Alpha 6 subunit of the 20S Core Protease of the proteasome | *α-6SU* | Nta.1081 | AACCAGTACGACACCGACGTAA | 59 | 145 | Pierella Karlusich *et al.*, 2017 |
| | | | CCTTGTTAACGCACGCGAGAAT | 59 | | |
| *N.tabacum* Elongation factor 1-α | *EF-1α* | AF120093.1 | TTCAGGAGCATGCGTCAAACTG | 55.9 | 100 | Schmidt and Delaney, 2010 |
| | | | TCTTCTTCTGAGCAGCCTTGGT | 55.6 | | |
| Putative *N.tabacum* 26S proteasome regulatory subunit 4 homolog A | *RPT2a* | XM_016599570.1 | TCAAGAAGAAAGAGGGAGTG | 54.6 | 141 | This study |
| | | | GAAGATCAGAAGGCAACATAG | 54.2 | | |
| Putative *N.tabacum* 26S proteasome regulatory subunit 6A homolog | *RPT5a* | XM_016627710.1 | CTGGTCGATTGGATCGTAAG | 58.2 | 102 | This study |
| | | | CGTCTGGGTTAACATTCATC | 55.5 | | |
